# Identification and genetic validation of drug-comorbidity interactions in type 2 diabetes using data-driven disease trajectory analysis

**DOI:** 10.1101/814392

**Authors:** Osamu Ichikawa, Benjamin S. Glicksberg, Noga Minsky, Hao-Chih Lee, Shuyu D. Li, Rong Chen, Li Li, Joel T. Dudley

## Abstract

Given various risk profiles, manifestations, comorbidities, and outcomes for individuals with Type 2 diabetes mellitus (T2D), a one-size-fits-all approach towards treatment and management is inadequate, and there is a clear need for personalized medications to reduce rates of complications and comorbidities. Here, we leveraged comprehensive Electronic Medical Records (EMR) to identify associations between T2D comorbidities and medications and evaluated their biological determinants using EMR-linked genetic information. We discovered clinically novel associations supported by the previous laboratory studies; e.g. 5-hydroxytryptamine 3 receptor antagonists, ondansetron and granisetron, are protective against cognitive disorders, and clopidogrel is protective against retinopathy. Furthermore, potentially novel associations were validated by genetic analysis; e.g. association between gabapentin and cognitive disorders was supported via variants of its target genes, *GRIN1* and *CACNA2D2*. These results of the current study open the door for optimizing treatment combinations to mitigate risks and for individualizing therapy based on T2D subtype risk profiles.

---

Type 2 diabetes mellitus (T2D) is a heterogeneous, multifactorial disease with a global diabetes prevalence that has more than doubled since 1980, rising from 4.3% to 9% in the adult population ^1^. Associated with psychological and physical distress to both patients and caregivers, diabetes is a considerable determinant of disability and remains the leading cause of vision loss, kidney failure, and non-traumatic lower limb amputations. Moreover, diabetes more than doubles the risk of the myocardial infarction and cerebrovascular accidents ^2, 3^. According to the National Health and Nutrition Examination Survey (NHANES) the prevalence of diabetes in the United States is over 12% ^4^, and as such, diabetes is associated with a total direct estimated medical cost of $237 billion in 2017 ^5, 6^. At the time of diagnosis 86% of patients with T2D are affected by chronic comorbid conditions ^7^, and this figure rises over the natural course of disease, compounding the burden on patients, providers, and the economy. There is tremendous need to advance the way in which T2D is treated in order to further reduce rates of complications and co-morbidities.

Given various risk profiles, manifestations, comorbidities, and outcomes for individuals with T2D, a one-size-fits-all approach towards treatment and management is inadequate. There have been attempts to disentangle determinants of poor patient outcomes, as well as efforts to develop more personalized treatment recommendations ^8^. One clear path forward is to model treatment recommendations based on disease profiles, since many conditions either co-manifest with or result from T2D. A previous study used Electronic Medical Records (EMR) to analyze patient characteristics and make treatments recommendations which improved diabetes control, measured by glycated hemoglobin A_1C_ (HbA1c), beyond the standard of care ^9^. Another study sought to evaluate the molecular effects of various medications on diabetes comorbidities using a systems biology approach ^10^.

In our previous work, we characterized the heterogeneous patient landscape of T2D into three distinct subtypes using clinical and genomic data from an EMR-paired biobank ^11^. We stratified patients using a topology-based, patient–patient network derived from demographic and clinical features such as disease diagnoses. We then layered on genetic information in the form of single nucleotide polymorphisms (SNPs) and characterized each subgroup according to enrichment of these features. For instance, subtype 1 was characterized by diabetic retinopathy and diabetic nephropathy; subtype 2 was enriched for lung cancer malignancy and cardiovascular diseases; and subtype 3 was associated most strongly with cardiovascular diseases and neurological diseases. Additionally, a recent study ^12^ discovered five replicable clusters of patients with T2D, representing different patient characteristics and risk of diabetic complications, based on a data-driven cluster analysis on six relevant variables. Taken together, the findings of those studies suggested that there might be subtypes of T2D that have different genetic and phenotypic characteristics, as well as risk factors for developing certain subsequent conditions.

As a data-driven analysis of a clinical population identified new T2D subtypes, we hypothesize that this approach can be applied to improve our knowledge regarding the association of medications with T2D related comorbidities and complications. In this study, we sought to push beyond our previous work to identify specific treatment strategies that are personalized according to T2D phenotypic patient profiles. Using the EMR of Mount Sinai Hospital, we investigated whether medications conferred a protective effect against, or the increased the risk of, subsequent comorbidities in all T2D patients (Fig. 1). Our results identified previously reported associations between medications and specific disease outcomes, and also uncovered potentially novel connections between medications and T2D comorbidities, some of which is supported by laboratory and animal data. We also evaluated the additive effects of medication pairs to identify potentially beneficial treatment regimens. In addition, we assessed relationships between predicted protective medications and disease comorbidities through a genetic association analysis of T2D patients in the Mount Sinai Bio*Me* biobank cohort, a subset of EMR patients. The results of the current study open the door for optimizing treatment combinations to mitigate risks and for individualizing therapy based on T2D subtype risk profiles.

**Fig. 1.**
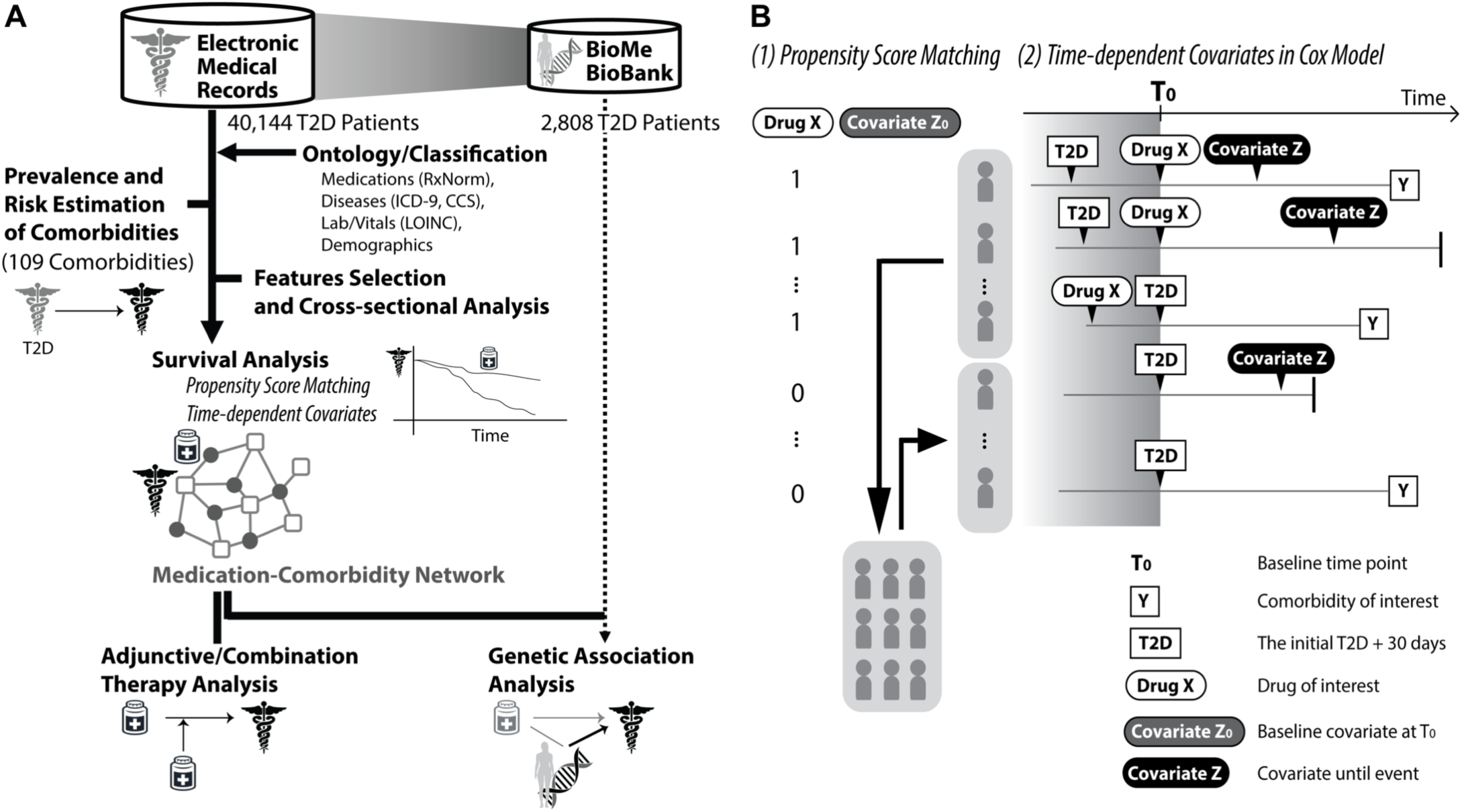
Schematic work flow of our study design and methods. (A) Workflow of current study. (B) Diagram of survival analysis with time dependent covariates after propensity score matching. The baseline time point, T_0_, was defined as the first prescription day of the targeted medication or 30 days after the initial T2D diagnosis date, whichever was later (see Supplementary Methods).

## Results

### Prevalence and risk estimation of comorbidities following T2D diagnosis

We began by assessing the prevalence and temporal ordering of the associations between T2D and other diseases (Fig. 1A). In particular, we sought to determine whether a given comorbidity tended to occur after diagnosis of T2D. We restricted our analysis to diseases with reported dates and only included the first reported encounter of a diagnosis, yielding 874,553 person–disease pairs, and analyzed their prevalence and temporal ordering at the patient level.

First, we calculated the prevalence of subsequent diseases based on the Kaplan–Meier product– limit estimator, in units of 1 year (Fig. 2A and Supplementary Table S1). In our cohort, the prevalence 10 years after T2D diagnosis was highest for ‘Hypertension’ (72% of cases), followed by ‘Disorder of lipid metabolism’ (63%). The macrovascular complication ‘Coronary atherosclerosis and other heart disease’ was had a prevalence of 46%, and the microvascular complication, ‘Chronic kidney disease’ was also quite common (34%). These results are consistent with previous studies demonstrating rates >67% for hypertension ^7^, 77% for ‘Disorder of lipid metabolism’ ^13^, 45–51% for ‘Coronary atherosclerosis and other heart disease’ ^14, 15^, and 24–40% for ‘Chronic kidney disease’ ^13, 16^, suggesting that our EMR data reflect a representative cohort of the US population. Ninety-six out of 219 diseases exhibited prevalence greater than 10% prevalence within 10 years after T2D diagnosis, and these were analyzed below. In addition, we analyzed all 13 neoplasms, because neoplasms are enriched in one of the T2D subtypes identified in our previous published study ^17^. 109 T2D comorbidities were analyzed in total.

**Fig. 2.**
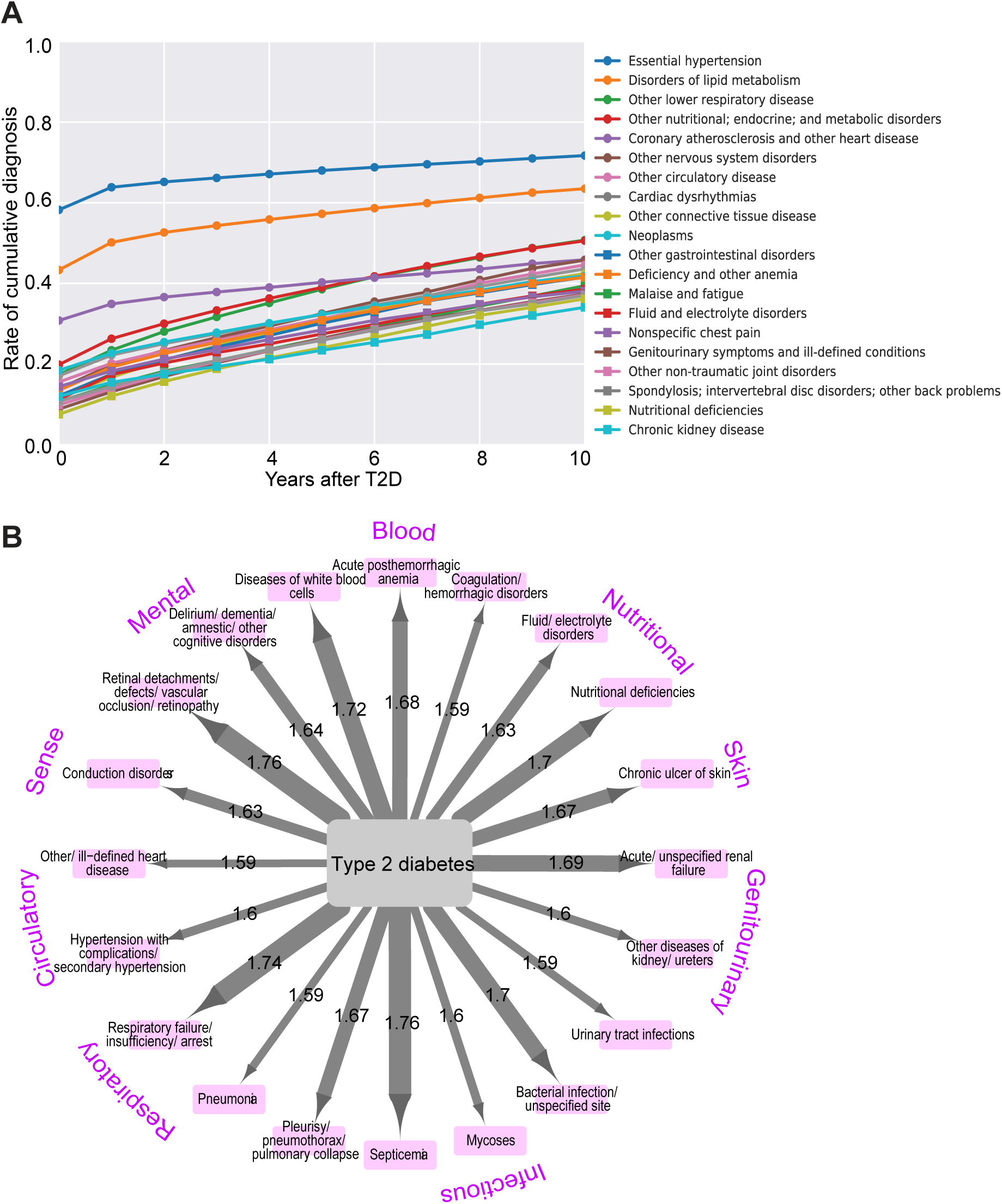
Prevalence and increased risk after T2D. (A) The 20 most prevalent diseases in our T2D cohort. The color coding of diseases is shown in the right of the plot. (B) The 20 diseases with the greatest increase in risk after T2D diagnosis (adjusted binomial p values < 0.05, using the Bonferroni method). The values on the edge and line thickness indicate relative risks. A complete list of prevalence and increased risk is provided in Supplementary Table S1.

**Table 1.** Protective or risk medications for the selected comorbidities.

| Comorbidity | Medication | HR cox | 95% CI | Adjusted<br>P value | Reference (PMID) |
| --- | --- | --- | --- | --- | --- |
| Retinal detachments; defects;<br>vascular occlusion; and<br>retinopathy | Psyllium | 0.47 | 0.26-0.83 | 3.35E-02 |  |
|  | Hydroxyzine | 0.52 | 0.33-0.81 | 1.66E-02 |  |
|  | Aluminum (IN<sup>a</sup>) | 0.52 | 0.32-0.85 | 3.23E-02 |  |
|  | Naproxen | 0.70 | 0.54-0.91 | 2.74E-02 |  |
|  | Lidocaine | 0.71 | 0.59-0.87 | 4.89E-03 |  |
|  | Nystatin | 0.72 | 0.57-0.92 | 2.67E-02 |  |
|  | Calcium | 0.73 | 0.57-0.92 | 3.20E-02 |  |
|  | Ibuprofen | 0.73 | 0.6-0.88 | 7.66E-03 |  |
|  | Clopidogrel | 0.80 | 0.68-0.94 | 2.54E-02 |  |
| Chronic kidney disease | Levetiracetam | 0.53 | 0.37-0.76 | 3.95E-03 |  |
|  | Metformin<sup>b</sup> | 0.63 | 0.63-0.63 | 0.00E+00 | 22641582, 21325870 |
|  | Sodium phosphate | 0.66 | 0.51-0.86 | 1.11E-02 |  |
|  | Ondansetron | 0.68 | 0.6-0.78 | 1.86E-06 |  |
|  | Naproxen | 0.69 | 0.54-0.9 | 2.38E-02 |  |
|  | Acetaminophen | 0.72 | 0.64-0.81 | 1.13E-06 |  |
|  | Hydrocortisone | 0.76 | 0.65-0.9 | 8.91E-03 | 28415636 |
| Coronary atherosclerosis and<br>other heart disease | Atovaquone | 0.22 | 0.1-0.52 | 3.76E-03 |  |
|  | Pseudoephedrine | 0.34 | 0.21-0.55 | 1.90E-04 |  |
|  | Timolol | 0.48 | 0.28-0.81 | 2.33E-02 | 15819591 |
|  | Diclofenac | 0.60 | 0.42-0.86 | 2.28E-02 |  |
|  | Potassium<br>phosphate | 0.60 | 0.45-0.81 | 5.52E-03 |  |
|  | Sulfamethoxazole | 0.62 | 0.48-0.81 | 3.95E-03 |  |
|  | Iron | 0.65 | 0.51-0.82 | 2.93E-03 | 18294483 |
|  | Cholecalciferol | 0.66 | 0.48-0.91 | 3.80E-02 | 27499590 |
|  | Hydromorphone | 0.74 | 0.6-0.92 | 2.55E-02 |  |
|  | Metformin<sup>b</sup> | 0.79 | 0.69-0.89 | 1.72E-03 | 28476766 |
| Acute cerebrovascular disease | Succinylcholine | 0.53 | 0.39-0.71 | 4.95E-04 |  |
|  | Repaglinide<sup>b</sup> | 0.54 | 0.37-0.8 | 1.02E-02 |  |
|  | Ondansetron | 0.59 | 0.47-0.75 | 1.84E-04 |  |
|  | Bacitracin | 0.61 | 0.42-0.89 | 3.60E-02 |  |
|  | Sitagliptin<sup>b</sup> | 0.65 | 0.46-0.92 | 4.67E-02 | 26687195 |
| Neoplasms excl. Benign | Betamethasone | 0.53 | 0.37-0.77 | 6.01E-03 |  |
|  | Aspirin | 0.57 | 0.57-0.57 | 0.00E+00 | 26940135 |
|  | Ketorolac | 0.65 | 0.53-0.79 | 4.78E-04 | 26071482 |
|  | Bacitracin | 0.67 | 0.51-0.87 | 1.56E-02 |  |
|  | Azithromycin | 0.72 | 0.62-0.84 | 4.11E-04 |  |
|  | Vancomycin | 0.73 | 0.61-0.88 | 4.88E-03 |  |
|  | Hydralazine | 0.74 | 0.61-0.89 | 8.55E-03 | 28277033 |
|  | Oxycodone | 0.79 | 0.7-0.9 | 3.39E-03 |  |
|  | Amoxicillin | 0.80 | 0.67-0.95 | 3.30E-02 | 28529929 |
|  | Lidocaine | 0.80 | 0.68-0.95 | 2.94E-02 |  |
| Delirium, dementia, and amnestic and other cognitive disorders | Granisetron | 0.44 | 0.22-0.85 | 4.48E-02 |  |
|  | Nifedipine | 0.54 | 0.42-0.7 | 8.29E-05 | 26221415 |
|  | Hydrocortisone | 0.65 | 0.53-0.81 | 1.64E-03 |  |
|  | Cefazolin | 0.68 | 0.58-0.8 | 3.42E-05 |  |
|  | Atorvastatin | 0.69 | 0.58-0.82 | 4.10E-04 | <sup>c</sup> 28212683, 27440840 |
|  | Ondansetron | 0.71 | 0.59-0.84 | 1.12E-03 |  |
|  | Hydromorphone | 0.72 | 0.58-0.89 | 1.23E-02 |  |
|  | Metformin<sup>b</sup> | 0.74 | 0.65-0.85 | 1.95E-04 | <sup>c</sup> 24577463, 28583443 |
|  | Gabapentin | 0.74 | 0.62-0.89 | 6.80E-03 |  |
|  | Phenylephrine | 0.76 | 0.64-0.92 | 1.87E-02 |  |
| Nutritional deficiencies | Psyllium | 0.59 | 0.44-0.8 | 3.68E-03 |  |
|  | Bupivacaine | 0.60 | 0.5-0.71 | 1.67E-07 |  |
|  | Gemfibrozil | 0.67 | 0.48-0.93 | 4.59E-02 |  |
|  | Acetaminophen | 0.67 | 0.61-0.74 | 6.28E-13 |  |
|  | Chlorhexidine | 0.67 | 0.5-0.91 | 3.09E-02 |  |
|  | Aspirin | 0.69 | 0.63-0.75 | 5.98E-14 |  |
|  | Atovaquone | 0.69 | 0.52-0.93 | 4.21E-02 |  |
|  | Dextromethorphan | 0.71 | 0.56-0.9 | 2.07E-02 |  |
|  | Tiotropium | 0.74 | 0.59-0.92 | 2.84E-02 |  |
|  | Allopurinol | 0.79 | 0.67-0.93 | 1.66E-02 |  |
|  | Clopidogrel | 0.79 | 0.71-0.88 | 1.98E-04 |  |

Table 1 (B)
| Comorbidity | Medication | HR cox | 95% CI | Adjusted P value | Reference (PMID) |
| --- | --- | --- | --- | --- | --- |
| Retinal detachments; defects; vascular occlusion; and retinopathy | Aprepitant | 4.81 | 2.4-9.65 | 1.84E-04 |  |
|  | HYLAN G-F 20 | 4.37 | 2.29-8.32 | 1.44E-04 |  |
|  | Prednisolone | 2.99 | 2.14-4.18 | 1.70E-08 |  |
|  | Desloratadine | 2.18 | 1.15-4.11 | 4.72E-02 |  |
|  | Timolol | 1.76 | 1.3-2.38 | 2.28E-03 |  |
|  | Repaglinide<sup>b</sup> | 1.47 | 1.18-1.82 | 3.86E-03 |  |
|  | Insulin<sup>b</sup> | 1.36 | 1.17-1.58 | 7.48E-04 |  |
|  | Lisinopril | 1.36 | 1.2-1.54 | 3.89E-05 |  |
|  | Glipizide | 1.20 | 1.04-1.38 | 3.66E-02 |  |
| Chronic kidney disease | Indigotindisulfonate | 2.65 | 1.55-4.55 | 3.26E-03 |  |
|  | Omega-3 Acid Ethyl Esters (USP) | 2.48 | 1.41-4.36 | 9.23E-03 |  |
|  | Calcitriol | 1.85 | 1.38-2.49 | 6.93E-04 |  |
|  | Pyridoxine | 1.64 | 1.15-2.34 | 2.43E-02 |  |
|  | Nifedipine | 1.52 | 1.3-1.78 | 9.49E-06 |  |
|  | Pseudoephedrine | 1.43 | 1.08-1.91 | 4.21E-02 |  |
|  | Prochlorperazine | 1.41 | 1.09-1.82 | 3.27E-02 |  |
|  | Insulin<sup>b</sup> | 1.40 | 1.25-1.58 | 1.05E-06 |  |
|  | Labetalol | 1.36 | 1.19-1.55 | 1.48E-04 |  |
|  | Ferrous sulfate | 1.30 | 1.14-1.48 | 1.15E-03 |  |
|  | Carvedilol | 1.18 | 1.04-1.34 | 3.24E-02 |  |
| Coronary atherosclerosis and other heart disease | Colchicine | 1.71 | 1.33-2.18 | 3.76E-04 |  |
|  | Insulin<sup>b</sup> | 1.19 | 1.03-1.37 | 4.89E-02 | <sup>c</sup> 28958751, 22686416 |
| Acute cerebrovascular disease | Phenytoin | 2.08 | 1.44-3.01 | 1.13E-03 |  |
| Neoplasms excl. Benign | Cosyntropin | 2.62 | 1.55-4.43 | 2.91E-03 |  |
|  | Trimethobenzamide | 2.10 | 1.27-3.48 | 1.77E-02 |  |
|  | Cetylpyridinium | 1.93 | 1.14-3.28 | 4.47E-02 |  |
|  | Nadolol | 1.91 | 1.38-2.64 | 1.19E-03 |  |
|  | Tenofovir | 1.85 | 1.3-2.62 | 4.61E-03 |  |
|  | Sodium sulfate | 1.46 | 1.09-1.94 | 3.39E-02 |  |
| Delirium, dementia, and amnestic and other cognitive disorders | Iohexol | 2.25 | 1.32-3.83 | 1.41E-02 |  |
|  | Bupropion | 1.59 | 1.22-2.08 | 4.32E-03 |  |
|  | Repaglinide<sup>b</sup> | 1.37 | 1.11-1.69 | 1.49E-02 |  |
|  | Heparin | 1.33 | 1.11-1.59 | 1.06E-02 |  |
| Nutritional deficiencies | HYLAN G-F 20 | 2.51 | 1.32-4.74 | 2.00E-02 |  |
|  | Ritonavir | 2.36 | 1.86-2.99 | 3.35E-10 | 26633015 |
|  | Thiopental | 2.11 | 1.19-3.76 | 3.54E-02 |  |
|  | Tretinoin | 1.95 | 1.19-3.19 | 2.81E-02 |  |
|  | Methotrexate | 1.60 | 1.18-2.18 | 1.32E-02 | 9517760 |
|  | Nateglinide<sup>b</sup> | 1.52 | 1.13-2.05 | 2.29E-02 |  |
|  | Dobutamine | 1.52 | 1.08-2.14 | 4.82E-02 |  |
|  | Fexofenadine | 1.47 | 1.2-1.79 | 1.67E-03 |  |
|  | Ramipril | 1.26 | 1.09-1.45 | 9.73E-03 |  |
|  | Homatropine | 1.26 | 1.07-1.48 | 2.49E-02 |  |
|  | Glyburide<sup>b</sup> | 1.24 | 1.06-1.44 | 2.33E-02 | 1070515 |
|  | Repaglinide<sup>b</sup> | 1.22 | 1.06-1.4 | 2.60E-02 |  |
|  | Metformin<sup>b</sup> | 1.14 | 1.06-1.22 | 2.31E-03 | 22179958 |

(A) Protective medications for the selected comorbidities (adjusted $p < 0.05$ , using Benjamini-Hochberg method). All comorbidities are listed in Supplementary Tables S4. (B) Risk medications for the selected comorbidities (adjusted $p < 0.05$ , using Benjamini-Hochberg method). All comorbidities are available in Supplementary Tables S5.
<sup>a</sup> Ingredient.
<sup>b</sup> Medications for T2D.
<sup>c</sup> Multiple published studies showed inconsistent results.

The increase in risk of each comorbidity after T2D diagnosis was quantified based on the timing of their occurrence in individual patients. Overall, out of the 109 T2D comorbidities analyzed, 93 diseases were significantly more likely to occur after T2D diagnosis (adjusted p < 0.05; Fig. 2B, Supplementary Table S1, and Supplementary Results).

### Candidate medications associated with each T2D comorbidity

Fifteen medications specifically indicated for T2D (T2D medications) were associated with at least one T2D comorbidity, with 79 protective and 198 risk associations (Fig. 3), in the cross-sectional analysis with adaptive LASSO (Supplementary Results). For example, insulin was associated with protection against 11 comorbidities and risk for 33 comorbidities, and metformin was associated with protection against 12 comorbidities and risk for 14 comorbidities. Rosiglitazone had the largest number of risk associations, including circulatory diseases. This is consistent with the previously reported risk of cardiovascular disease and death, which led to rosiglitazone being withdrawn from European and other markets ^18, 19^.

**Fig. 3.**
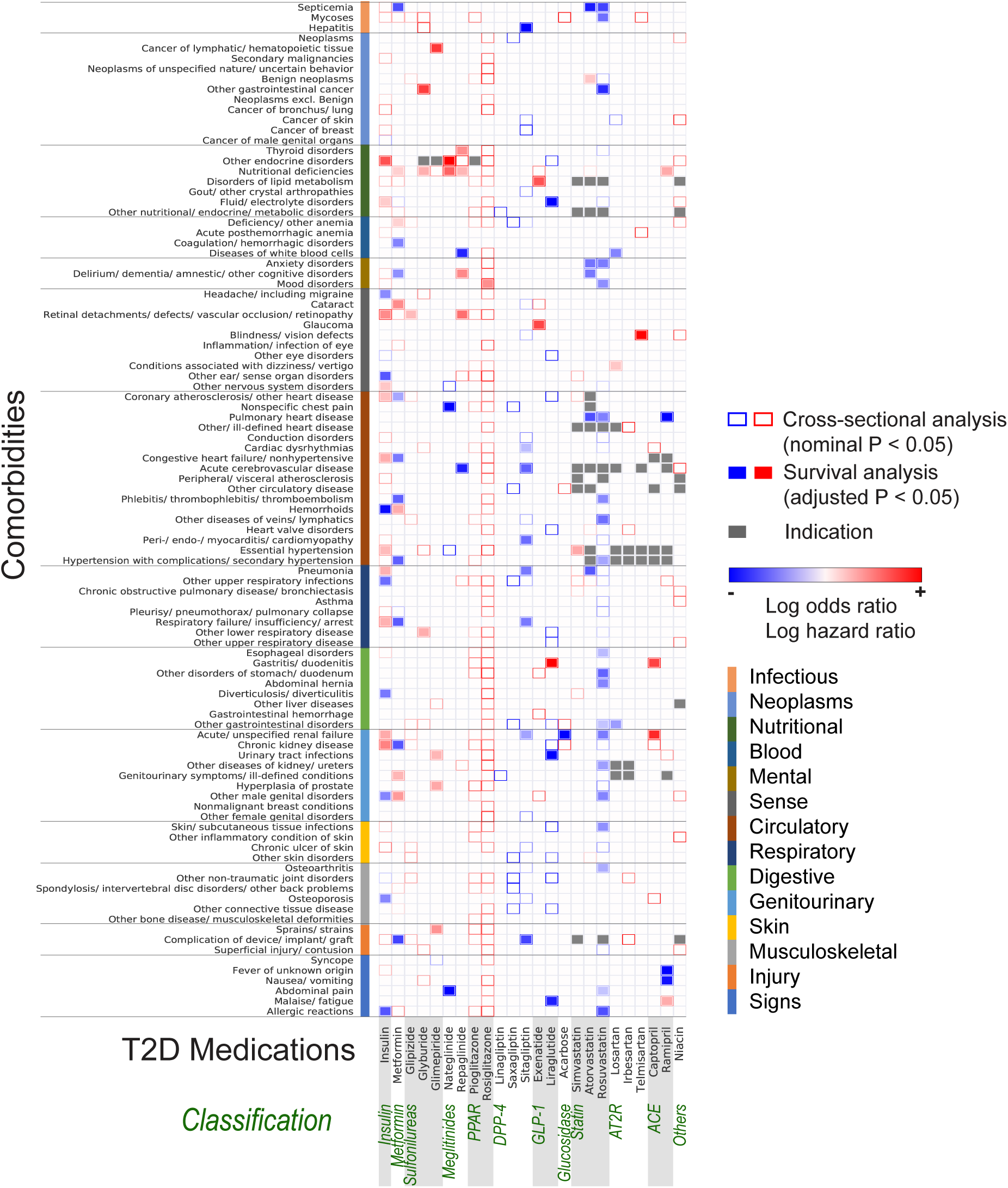
Heatmap of associations between T2D disease comorbidity and T2D /T2D-related medication. Rectangle border color represents odds ratio (OR) by the cross-sectional analysis using logistic regression (nominal p values < 0.05), and rectangle fill color represents hazard ratio (HR) by the survival analysis using time-dependent covariate Cox regression after propensity score matching (adjusted p values < 0.05, using the Benjamini-Hochberg method). The protective associations (OR < 1.0 or HR < 1.0) are displayed in blue, and the risk associations (OR > 1.0 or HR > 1.0) are displayed in red. The drug indications are colored gray.

Nine T2D-related medications, which are not necessarily indicated for T2D, but are often used for treating or managing T2D co-morbidities (see Supplementary Methods), were associated with at least one disease, including three statins and five antihypertensive medications (three angiotensin II receptor [AT2R] blockers, two angiotensin-converting-enzyme [ACE] inhibitors, and niacin) (Fig. 3). These medications had 48 protective associations and 42 risk associations. Two high potency statins, atorvastatin and rosuvastatin, exhibited a wide variety of protective effects but few risks for T2D comorbidities, supporting that the use of these medications is a key factor in preventing and managing comorbidities related to lipid metabolism ^20^.

### T2D and T2D-related medications predicted to modulate the onset of various comorbidities

Among the candidate medications associated with each T2D comorbidity in the cross-sectional analysis, we assessed the timing effect of medications on disease onset by survival analysis with time-dependent covariates in propensity score–matched cohorts (Fig. 1B). Among T2D medications, indicated directly for diabetes treatment (see Supplementary Methods), and T2D-related medications, 64 out of 127 protective associations and 43 of 240 risk associations were also identified by the survival analysis (Fig. 3 and Supplementary Table S3). The analysis identified 30 medication-disease associations described previously, involving the medications commonly prescribed in patients with T2D, such as metformin, insulin, and statins. These associations include, for instance, atorvastatin and ‘Pulmonary heart disease’ (HR=0.62, adjusted p < 0.0001) ^21^, sitagliptin and ‘Hepatitis’ (HR=0.43, adjusted p=0.0060) ^22^, metformin and ‘Nutritional deficiency’ (HR=1.1, adjusted p=0.0023) ^23^, and nateglinide and ‘Other endocrine disorders’, which includes hypoglycemia (HR=3.7, adjusted p< 0.0001) ^24^ (Supplementary Table S3). Some associations were controversial due to inconsistent results in published studies. Among them, five medications had protective associations in our study, including metformin with ‘Delirium, dementia, and amnestic and other cognitive disorders’ (HR=0.74, adjusted p=0.00020) ^25, 26^; atorvastatin with ‘Delirium, dementia, and amnestic and other cognitive’ (HR=0.69, adjusted p=0.00041) ^27, 28^; rosuvastatin with ‘Esophageal disorders’ (HR=0.84, adjusted p=0.040) ^29, 30^; and sitagliptin with ‘Acute and unspecified renal failure’ (HR=0.78, adjusted p=0.027) ^31^ and ‘Cardiac dysrhythmias’ (HR=0.84, adjusted p=0.047) ^32, 33^. Lastly, we identified 40 protective associations which have not previously been described. These include insulin and ‘Osteoporosis’ (HR=0.72, adjusted p=0.0017), liraglutide and ‘Urinary tract infections’ (HR=0.46, adjusted p=0.048), and nateglinide and ‘Abdominal pain’ (HR=0.38, adjusted p=0.013). In addition, we identified 30 new risk associations, including exenatide and ‘Glaucoma’ (HR=1.6, adjusted p=0.046), metformin and ‘Cataract’ (HR=1.36, adjusted p < 0.0001), and nateglinide and ‘Nutritional deficiencies’ (HR=1.52, adjusted p=0.023).

### Non-T2D-related medications predicted to modulate the onset of well-known T2D complications

Two hundred thirty one medications, including non-T2D-related medications, exhibited protective effects for at least one of the 104 T2D comorbidities (Fig. 4 and Supplementary Table S4), and 270 medications increased the risk for at least one comorbidity (Fig. 4 and Supplementary Table S5). Table 1 summarizes seven comorbidities, including two major microvascular complications, ‘Retinal detachments; defects; vascular occlusion; and retinopathy’ and ‘Chronic kidney disease’; two major macrovascular complications ‘Coronary atherosclerosis and other heart disease’ and ‘Acute cerebrovascular disease’; and three comorbidities that were enriched in particular T2D subtypes in our previously published study ^11^, ‘Neoplasm excl. benign’, ‘Delirium, dementia, and amnestic and other cognitive disorders’, and ‘Nutritional deficiencies’.

**Fig. 4.**
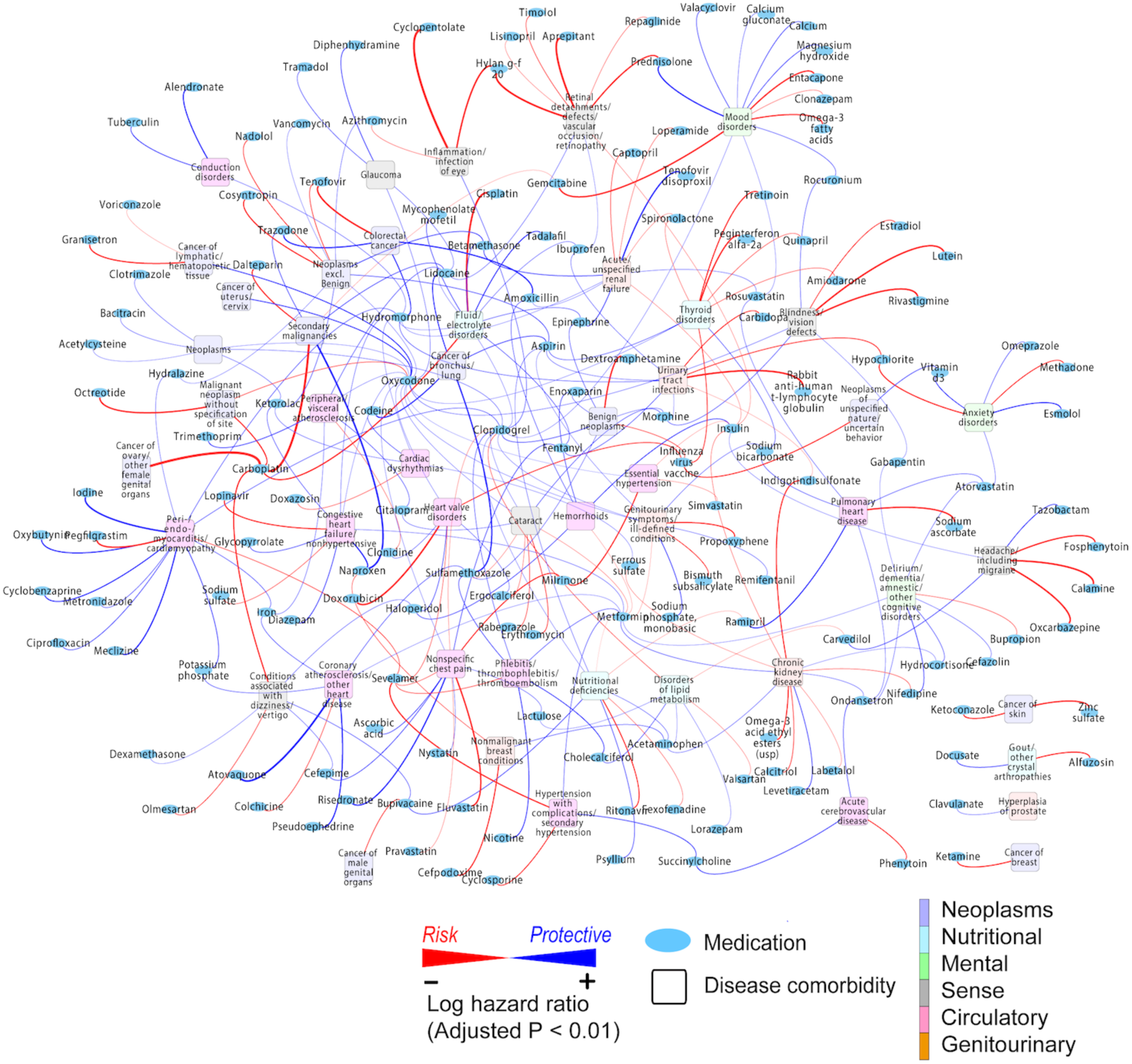
Network of T2D disease comorbidities and medications. This network includes all kinds of medications, including non-T2D medications, and associations with adjusted p values < 0.01, using the Benjamini-Hochberg method. For visualization purposes, we showed the diseases categorized into six Multi-Level CCS codes (Neoplasms, Nutritional, Mental, Sense, Circulatory, and Genitourinary), except for residual diseases beginning with “other” (e.g., “Other circulatory disease”). Blue ellipses represent medications, and squares represent disease comorbidities with the fill color corresponding to the disease categories by Multi-Level CCS codes. The protective associations (HR < 1.0) are displayed by blue lines, and the risk associations (HR > 1.0) are displayed by red lines. Line thickness represents the magnitude of HR. A complete list of all associations (adjusted p values < 0.05) is available in Supplementary Tables S4 and S5.

We identified medications whose use conferred protective effects against the development of specific comorbidities in T2D patients (Table 1A). Twelve medications were demonstrated to have protective associations which were previously described, including cholecalciferol and ‘Coronary atherosclerosis and other heart disease’ (HR=0.66, adjusted p=0.038) ^34^; aspirin and ‘Neoplasm excl. benign’ (HR=0.57, adjusted p<0.0001) ^35^; and nifedipine and ‘Delirium, dementia, and amnestic and other cognitive disorders’ (HR=0.54, adjusted p<0.0001) ^36^. Intriguingly, our analysis confirms protective medication effects previously demonstrated only in non-clinical laboratory studies. Two 5-hydroxytryptamine 3 (5-HT_3_) receptor antagonists (ondansetron and granisetron) and one alpha-adrenergic agonist (phenylephrine), exhibited protective effects against ‘Delirium, dementia, and amnestic and other cognitive disorders’ (HR=0.71, adjusted p=0.0011; HR=0.44, adjusted p=0.045; HR=0.76, adjusted p=0.019; respectively) ^37, 38^. Bacitracin, azithromycin, and lidocaine protected against ‘Neoplasm excl. benign’ (HR=0.67, adjusted p=0.016; HR=0.72, adjusted p=0.00041; HR=0.80, adjusted p=0.029; respectively) ^39–41^. Clopidogrel and hydroxyzine were protective against ‘Retinal detachments; defects; vascular occlusion; and retinopathy’ (HR=0.80, adjusted p=0.025; HR=0.52, adjusted p=0.017; respectively) ^42, 43^.

We identified a list of potentially novel protective associations (Table 1A). In regard to microvascular complications, seven medications were significantly associated with ‘Retinal detachments; defects; vascular occlusion; and retinopathy’, and five medications were related to ‘Chronic kidney disease’. In regard to macrovascular complications, six medications were associated with ‘Coronary atherosclerosis and other heart disease’, and four medications exhibited a protective effect on ‘Acute cerebrovascular disease’. Four medications, including gabapentin (HR=0.74, adjusted p=0.0068), one of the first-line medications for the treatment of neuropathic pain in diabetic neuropathy, exhibited protective effects against ‘Delirium, dementia, and amnestic and other cognitive disorders’. Three medications, vancomycin (HR=0.73, adjusted p=0.0049), betamethasone (HR=0.53, adjusted p=0.0060), and oxycodone (HR=0.79, adjusted p=0.0034), were associated with protection against ‘Neoplasm excl. benign’ and 11 medications were related to ‘Nutritional deficiencies’. A complete list of all disease categories is provided in Supplementary Table S4.

We also discovered associations between medications and increased risk for comorbidities (Table 1B). We corroborated two previously reported risk medications, except for T2D medications, to ‘Nutritional deficiencies’, which were methotrexate (HR=1.6, adjusted p=0.013) ^44^ and ritonavir (HR=2.4, adjusted p<0.0001) ^45^. In addition, we identified 9 potentially novel associations between medications and ‘Retinal detachments; defects; vascular occlusion; and retinopathy’ and 11 potentially novel associations between medications and ‘Chronic kidney disease’ (Table 1B). Colchicine was associated with increased risk of ‘Coronary atherosclerosis and other heart disease’ (HR=1.7, adjusted p=0.00038), and phenytoin with increased risk of ‘Acute cerebrovascular disease’ (HR=2.1, adjusted p=0.0011). Four medications were associated with risk of ‘Delirium, dementia, and amnestic and other cognitive disorders’. Six medications were associated with an increased risk of ‘Neoplasms excl. benign’. Nine medications were associated with ‘Nutritional deficiencies’. A complete list of all disease categories is available in Supplementary Table S5.

### Validation of the predicted comorbidity-medication relationship by genetic association analysis

We evaluated the associations between the predicted medications and seven disease comorbidities of interest (Table 1A) within the Mount Sinai Bio*Me* biobank cohort (see Methods). Specifically, we analyzed the associations between each comorbidity and SNPs mapped to the genes targeted by medications predicted to be protective.

We found that 286 of 4,912 SNPs were significantly associated with comorbidities, with nominal p values less than 0.05 (Table 2 and Supplementary Table S9). We identified 58 SNPs, corresponding to 17 unique drug targets, that were significantly enriched in patients diagnosed with ‘Retinal detachments; defects; vascular occlusion; and retinopathy’. These patients could, in theory, benefit from three corresponding medications: calcium, hydroxyzine, and ibuprofen, which demonstrated a protective effect. Twelve SNPs mapped to five genes targeted by metformin, naproxen, and ondansetron were associated with ‘Chronic kidney disease’. Additionally, 45 SNPs mapped to 18 genes connected to six medications were associated with ‘Coronary atherosclerosis and other heart disease’. Thirty-four SNPs mapped to 11 genes targeted by five medications were associated with ‘Acute cerebrovascular disease’. Twenty-five SNPs corresponding to six medications, specifically azithromycin, bacitracin, betamethasone, ketorolac, lidocaine, and oxycodone, were related to ‘Neoplasms excluding benign’. Seventy-seven SNPs mapped to 15 genes related to four medications, gabapentin, nifedipine, cefazolin, and phenylephrine, were associated with ‘Delirium, dementia, and amnestic and other cognitive disorders.’ Sixteen SNPs linked to four medications were related to ‘Nutritional deficiencies’. A complete list of each significant SNP-comorbidity association is provided in Supplementary Table S9.

**Table 2.**
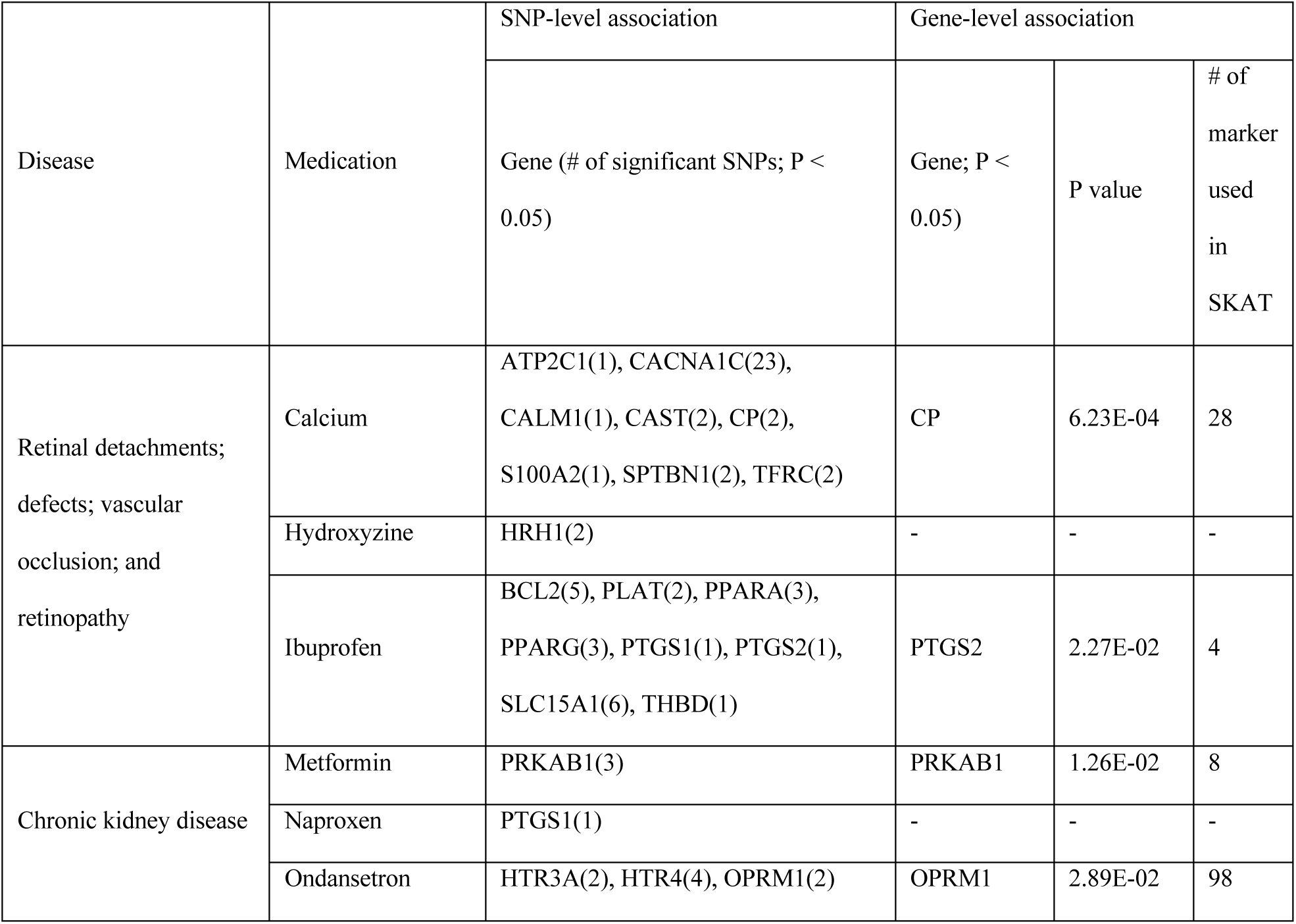
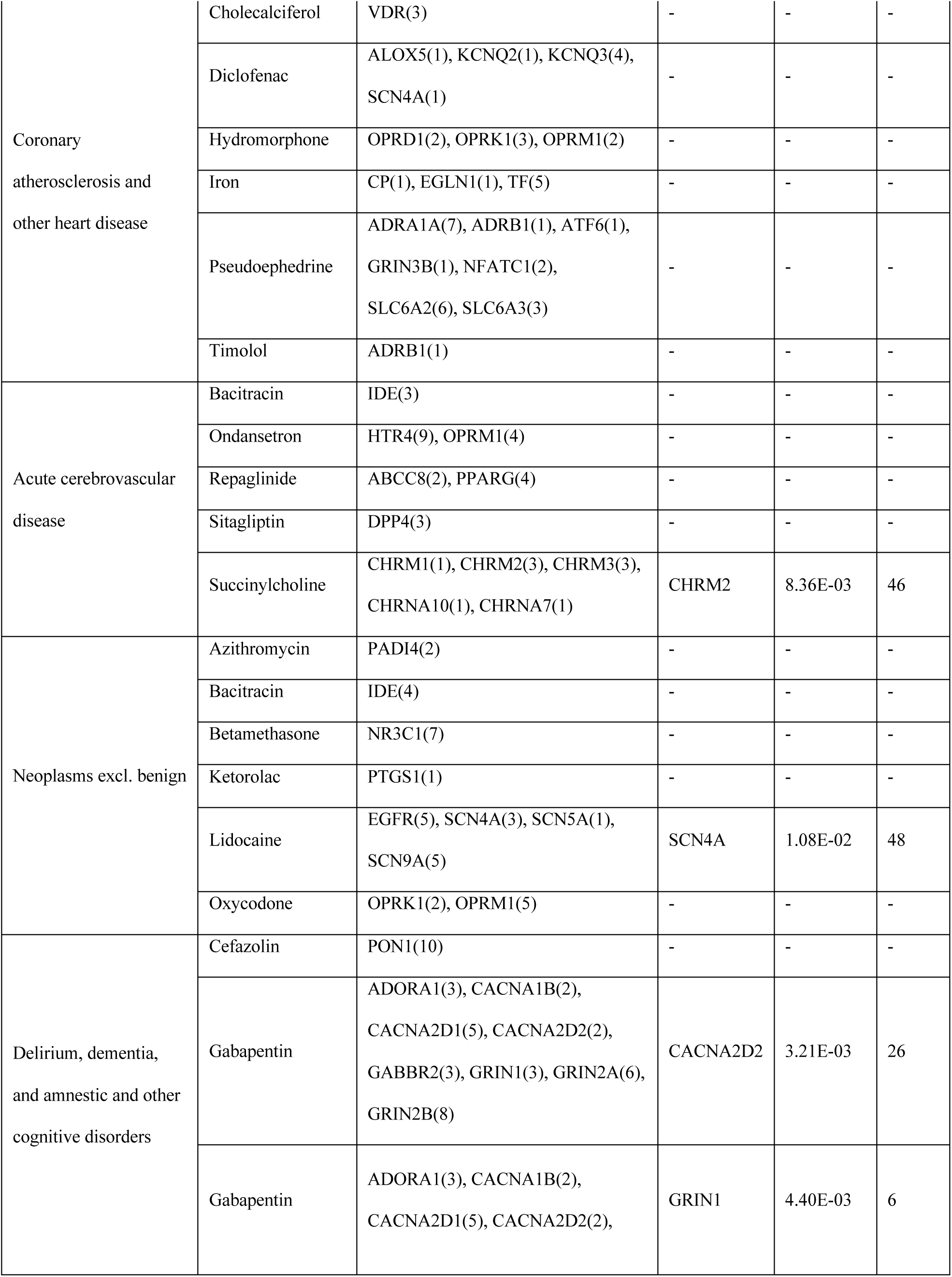
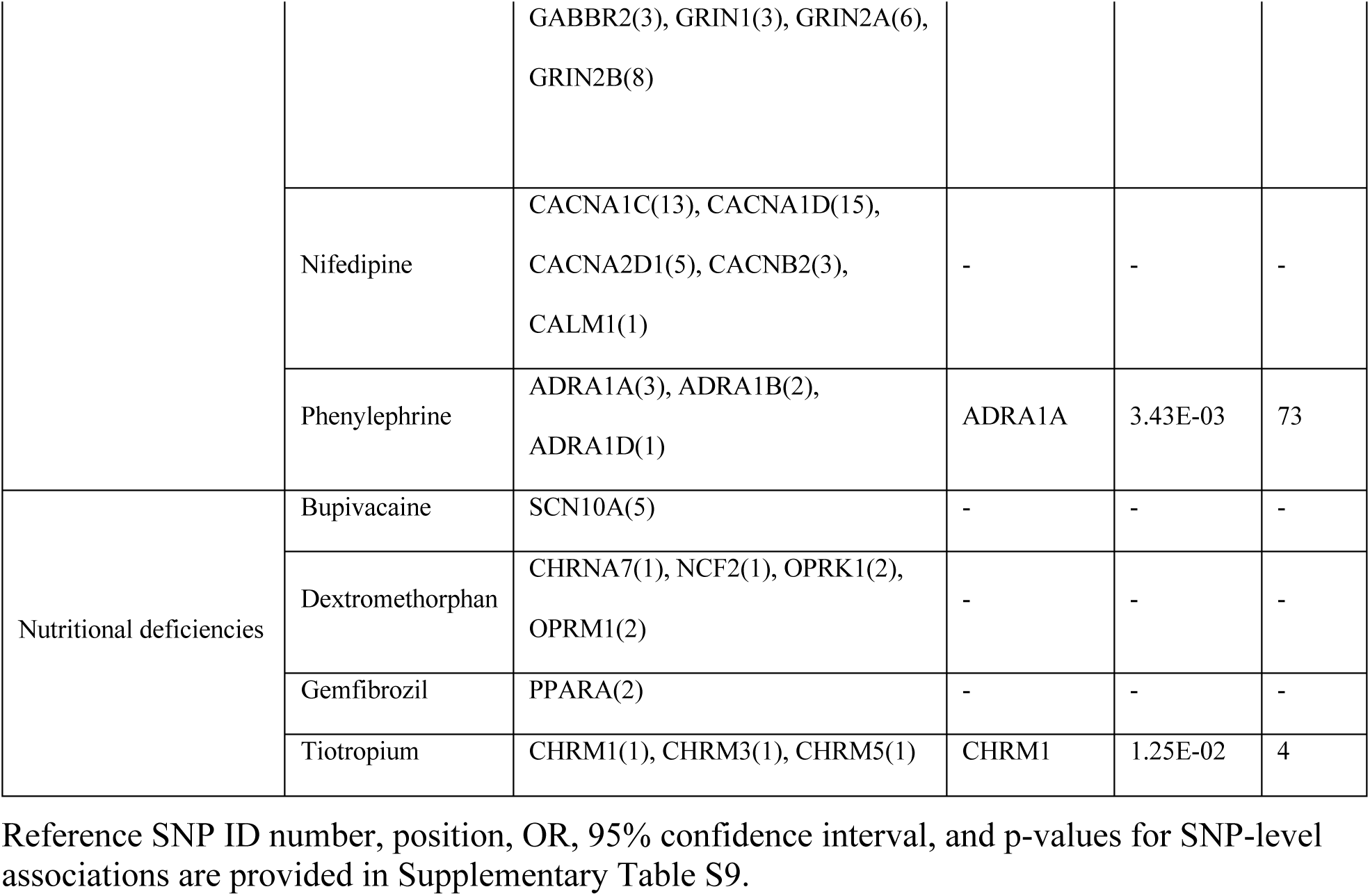
Summary of associations between T2D comorbidities and SNPs/genes related to predicted protective medications (p<0.05).

In the SNP set–comorbidity combinations, we gathered SNPs based on gene-level annotations. There were nine associations between the gene level and comorbidities, all of which were consistently identified within the SNP–comorbidity combinations described above (Table 2).

These genetic associations between drug-target genes/SNPs and comorbidities in the T2D patient cohort strongly support our findings regarding the protective associations between medications and comorbidities and suggest mechanisms of actions for each medication.

## Discussion

Because T2D is associated with various clinical manifestations and chronic comorbidities over time, the optimal therapeutic regimen evolves according to the changing disease profile of the individual patient. In this study, we developed the first data-driven deep phenotyping strategy for identifying the risk and benefits of medications based on the T2D comorbidity profiles. These T2D comorbidity profiles are based on our comprehensive large-scale EMR, which contains longitudinal clinical measurements, prescribed or administered medications, and disease diagnoses. Analyzing EMR enables us to explore longitudinal trajectory outcomes with more than 10 years of follow-up, increasing the likelihood of capturing long-term effects. We developed a systematic approach to estimate the association of medications, alone in combination, with future onset of comorbidities. Our approach could facilitate the precision of treatment regimens tailored to individual disease profiles.

Our analyses confirmed known associations between medications and diseases, and also uncovered potentially novel connections between medications and T2D comorbidities. Our approach began with evaluating the prevalence of comorbidities in T2D patients after the initial diagnosis of T2D. Next, statistical models were applied to assess the effects of medications on the risk of developing subsequent comorbidities. Lastly, medications and disease associations were assessed and explained by genetic variants using the T2D patient cohort in the genotyping biobank data from our Bio*Me* program_17._

Our analysis provides clinical evidence for associations previously demonstrated only in laboratory and animal studies. Two 5-hydroxytryptamine 3 (5-HT_3_) receptor antagonists, ondansetron and granisetron, consistently showed protective effects to ‘Delirium, dementia, and amnestic and other cognitive disorders’. Several studies on rats and nonhuman primates showed that ondansetron has positive effects on learning ^37^. A clinical trial of ondansetron ^46^ revealed no significant therapeutic benefit for Alzheimer’s disease (AD), but that study was conducted on mild/moderate AD patients who received a short-term treatment (24 weeks), rather than pre-AD patients in which the condition might be prevented. This result, in conjunction with other two studies with 5-HT reuptake inhibitors ^47, 48^, suggest that serotonergic strategies are not effective at improving cognition in patients with existing AD. Nevertheless, our analysis is the first to support a protective effect of 5-HT_3_ antagonists against cognitive disorders, the first such evidence to be obtained in human patients. Thus, our EMR-based studies could reveal the long-term effects of prognostic factors for pre-dementia patients, particularly patients with T2D. To our knowledge, we are the first to describe a protective association in humans between clopidogrel, classified as a thienopyridine, and ‘Retinal detachments; defects; vascular occlusion; and retinopathy’. A protective effect was established for ischemic retinopathy in a mouse model ^43^. A clinical trial using a randomized, double-masked, placebo-controlled design showed that another thienopyridine, ticlopidine, significantly slows the progression of nonproliferative diabetic retinopathy in insulin-treated diabetic patients ^49^, and our result supports a protective class effect. Additionally, our analysis supported the previous studies, which have reported that some antibiotics are associated with decreased risk for neoplasms (See Supplementary Discussion).

Genetic analysis of the T2D patient cohort in our biobank provided potential insights into the protective associations between medication and disease comorbidities through an inferred underlying mechanism of action (Table 2). The potentially novel association between ‘Delirium, dementia, and amnestic and other cognitive disorders’ and the antiepileptic gabapentin, a first-line medications for neuropathic pain in diabetic neuropathy, is supported by the relationship between this disease and its target genes, *GRIN1* and *CACNA2D2* (Table 2). *GRIN1* encodes NMDA (N-methyl-D-aspartate) receptor 1, a target gene of the Alzheimer’s disease drug, memantin. In addition, voltage-gated calcium channels, including CACNA2D2, are inhibited by gabapentin ^50^; calcium channel blockade attenuates amyloid-β-induced neuronal decline *in vitro* and is neuroprotective in animal models ^51^. Further potentially novel associations were discussed in Supplementary Discussion.

Although some of the medication–comorbidity associations we identified have been supported by previous preclinical studies, the genetic association results further strengthen and guide our interpretation of the potential mechanisms of action. The association between ‘Retinal detachments; defects; vascular occlusion; and retinopathy’ and the histamine H1 blocker hydroxyzine is supported by two SNPs mapped to *HRH1*. The protective effect of an H1 blocker against retinopathy is consistent with a previous study showing that H1 blockage is effective in reversing leakage of albumin from the blood across the retinal barrier, a typical phenotype of diabetic retinopathy in diabetic rats ^52^. We also describe an association between lidocaine and lower rates of ‘Neoplasm excluding benign’, and this observation is supported by studies which demonstrated that a clinical concentration of lidocaine inhibits epidermal growth factor receptor (EGFR) activity and suppresses EGF-induced in vitro cancer cell proliferation ^53, 54^.

Though several of the associations demonstrated in our analysis have been previously reported by patient-level studies, our genetic association analysis sheds light on potential mechanisms of drug action. For example, the protective effect of metformin against ‘Chronic kidney disease’ is supported by the association with *PRKAB1*, the gene coding AMP-activated protein kinase (*AMPK*), which is activated by metformin. *AMPK* is an energy sensor that plays a pivotal role in cellular homoeostasis. Deficiency in *AMPK* activity promotes epithelial-to-mesenchymal transition and renal cell apoptosis, eventually contributing to chronic kidney disease (CKD) ^55^. In addition, a meta-analysis of genome-wide association studies has identified a link between estimated glomerular filtration rate (eGFR) and another component of *AMPK*, encoded by *PRKAG2* ^56^. Although, due to a risk of lactic acidosis, metformin is currently contraindicated in patients with severe CKD (eGFR < 30 ml/min/1.73 m^2^), it has been suggested that the benefits of its use may outweigh possible risks in these patients ^57^. A novel *AMPK* activator, without the risk of lactic acidosis, could be a safer alternative to metformin in preventing progression of severe CKD.

In addition to identifying associations between medications and comorbidities, we analyzed whether medication pairing modified these associations (Supplementary Results and Supplementary Table S7). Some T2D medications were associated with nutritional deficiencies and this implies closer monitoring may be necessary with certain drugs. We observed novel associations between metformin and nateglinide with vitamin D deficiency, which is interesting because of data supporting that vitamin D supplementation may improve insulin sensitivity ^58^. Other novel associations identified include glyburide and vitamin B-complex deficiencies and repaglinide and protein-calorie malnutrition.

In our current study, we build upon our previous work to identify specific treatment strategies according to individual T2D disease profiles and drug-disease associations. In our preceding publication, we characterized the heterogeneous patient landscape of T2D into three distinct subtypes using clinical data from an EMR and verified the disease association in a genetics biobank ^11^. That work had suggested that there might be patterns of T2D with different genetic associated phenotypic characteristics, as well as risk factors for developing certain subsequent conditions. Combining this knowledge with our current observations regarding medication associations with comorbidities, a more personalized and precise approach toward medication regimens can be devised (Supplementary Discussion).

This study has its limitations. First, to define the patient cohort, we had to use ICD-9 codes instead of the well-validated eMERGE algorithm ^59^ because about 30% of the patients did not have lab test results for random/fasting glucose or HbA1c. To ensure high-quality phenotyping, however, we carefully ruled out type 1 diabetes mellitus, similarly to what is done by the eMERGE algorithm. Furthermore, the eMERGE algorithm was developed in 2012 and does not capture T2D medications approved by FDA after 2012, such as some DPP-4 inhibitors, GLP-1 agonists, insulin analogs, and any SGLT-2 inhibitors. Second, the first diagnosis of T2D may not always have been documented at our hospital, so we cannot ensure that a given patient was not been diagnosed previously elsewhere. By the same reasoning, it is possible that some diagnoses and prescription of medications documented at different institutions may have been missed. One possible solution and future direction is to incorporate data from insurance claims. In addition, the definition of outcome comorbidities by CCS Diagnoses terms would reduce the resolution of disease category coded by ICD-9. However, we could collect sufficient sample size and reduce the noise of underdiagnoses, including assignment of a similar but different ICD-9 code, which was observed in diabetic neuropathy. Finally, the results of this observational study need to be interpreted with caution and will require confirmation in randomized clinical trials that stratify treatment groups based on the status of disease comorbidities and other clinical features.

Our study is the first to investigate a comprehensive EMR linked to a biobank with the aim of discovering the risk of T2D comorbidities and revealing treatment strategies tailored to patient risk profiles. Using systematic feature selection with a regularization technique and survival analysis with time dependent covariate in propensity score matched cohort, we identified a list of medications that may influence the risk spectrum of T2D comorbidities. We analyzed medications based on ingredient level irrespectively on drug form, and the approach enabled us to identify the ingredients that associate with comorbidities, which have not been reported in literature. Future studies would be followed up for further investigation. Finally, both previously observed and novel associations between medications and comorbidities were validated by genetic variants analysis using the T2D patient cohort in our biobank. With further evidence, if these recommendations can successfully be implemented in the clinic, it would undoubtedly reduce the economic burden by prioritizing prescription of the right medication at the right time. Our medication–comorbidity model provides a real-world, data-driven approach for treatment stratifications that consider future development of comorbidities, with the ultimate goal of guiding precision medicine based on individual clinical phenotypes of patients with T2D.

## Methods

### Ethics statement

All data were fully anonymized. Ethical approval was provided by Program for the Protection of Human Subjects (PPHS) at Mount Sinai health system (HS#17-01058, IRB#17-02530). Waiver of informed consent was approved due to the purpose and scale of this study with retrospective and anonymous data only.

### Study design

The objective of our study was to develop a data-driven approach to identify the risk and benefits of medications based on the T2D comorbidity profiles for guiding precision medicine tailored to individual clinical phenotypes. We used comprehensive Electronic Medical Records (EMR) system of Mount Sinai Hospital (MSH) to identify associations between T2D comorbidities and medications. We analyzed a T2D cohort of 40,144 individuals with well-annotated patient information. Please refer to Supplementary Methods section for patient cohort, clinical sources and term standardization, and definition of outcomes. We applied a systematic feature selection using a regularization technique and survival analysis method with time dependent covariates in a propensity score matched cohort. The relations between medications and comorbidities were validated by genetic variants analysis using the T2D patient cohort in our EMR-linked biobank.

### Disease prevalence and disease pair temporal directionality

We calculated the disease prevalence based on the Kaplan–Meier product–limit estimator, with units of 1 year, and then we assessed disease-pair connectivity patterns for T2D comorbidities. Please refer to the Supplementary Methods section for detailed descriptions of the disease prevalence and disease pair temporal directionality analyses.

### Feature selection

We considered many disease variables coded by Single-Level Clinical Classifications Software (CCS) diagnoses, medication variables mapped to RxNorm ingredient code, and lab/vital variables based on LOINC codes (Supplementary Methods). Accordingly, we adopted a feature selection method, specifically a penalized logistic regression with adaptive LASSO (Eq. 1), to identify variables of the highest relevance associated with ensuing comorbidities following T2D disease diagnosis. First, we pre-filtered disease and medication variables to only include those with more than 20 patients, and lab/vital variables measured for more than 50% of patients, yielding a total of 1,255 features including 246 diseases, 899 medications, 107 lab/vitals, and four demographic features. Adaptive LASSO is an extension of traditional LASSO ^60^ that uses coefficient-specific weights ^61^. The adaptive LASSO estimator may achieve sparsity and selection consistency for the true model, i.e., correctly identifies the zero and nonzero parameters ^62^. Let *L_n_*(*β Y, X*) be the negative log-likelihood parametrized by β for a sample of size *n*. The adaptive LASSO estimator is defined as:

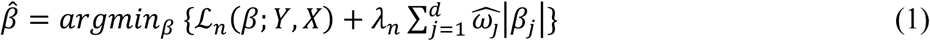

Where 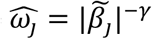 is a coefficient specific weights vector, and *λ_n_* is a regularization parameter. We set the positive constant γ as 1, according to Zou et al. ^61^, and obtained β̃ by the maximum likelihood estimate of Ridge regression. The *λ_n_* value for minimum AUC was chosen by 10-fold cross validation. We used the R package *glmnet* ^63^ for these penalized regressions.

### Logistic regression model

In the cross-sectional analysis, we used the odds ratio (OR) from logistic regression to quantify the magnitude of the risk of comorbidity associated with prescription of each medication (i.e., increased risk or protective effect). Please see the Supplementary Methods section for a detailed description of logistic regression. Features that were significantly associated with each of the outcome comorbid diseases with nominal p values less than 0.05 were used for the following propensity score matching and survival analysis.

### Survival analysis

We calculated hazard ratios using Cox proportional hazards models with propensity score matching of cases and controls. Propensity score matching is described in Supplementary Methods. Although many previous studies demonstrated that propensity score matching effectively suppresses confounder effects ^64, 65^, we further adjusted the time-dependent effects after the baseline time point using time-dependent covariates ^66^. We did so because our study focuses on the association between medications and the development of comorbid diseases, considering interventions after the baseline time point (Fig. 1B). For the time-dependent covariates, we used as many adaptive-LASSO-selected covariates as possible unless a given covariate did not violate the proportional hazards assumption.

Cox proportional hazard model with time-dependent covariates:

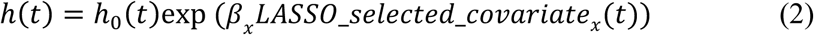

where *h*(*t*) is the expected hazard at time *t*, *h*_0_(*t*) is the baseline hazard, and

*LASSO_selected_covariate_x_(*t*)* is each of the time *t*-dependent binary variables selected by adaptive LASSO, such as medications, disease history, lab values, and vital signs.

We confirmed the proportional hazards assumption by calculating Schoenfeld residuals (Schoenfeld test p>0.05). We used the R packages *survival* and *survminer* for survival analysis. The p values derived from the Cox models were adjusted for multiple testing using the Benjamini and Hochberg false discovery rate method ^67^.

### SNP/gene and disease association analyses

We assessed our predicted associations between medications and above seven well-known T2D comorbidities ^11^ by performing a genetic association analysis. Specifically, we retrieved drug–gene relationship information from DrugBank version 5.0.11 ^68^, and then obtained genotyping data for all SNPs mapped to these genes in our cohort. We analyzed association a SNP–comorbidity basis as well as a gene–comorbidity basis by using the SNP-set Kernel Association Test (SKAT) method ^69^ (Supplementary Methods).

## Data availability

The data that support the findings of this study are available from the corresponding author upon reasonable request.

## Acknowledgements

We thank Mount Sinai Data Warehouse for data access and infrastructural support. The Bio*Me* Biobank was funded by The Charles Bronfman Institute for Personalized Medicine at the Icahn School of Medicine at Mount Sinai. We also thank Douglas Ruderfer and Eli Stahl for performing the original quality control of the Bio*Me* genotype data. This work was supported in part by a gift from the Harris Family Charitable Foundation (to J.T. D.), and grant from the NIH R01 DK098242. The funders had no role in study design, data collection and analysis, decision to publish, or preparation of the manuscript.

## Author Contributions

Study supervision: L.L. and J.T.D. Conceived and designed the study: O.I. and L.L. Analyzed EMRs: O.I., B.G., and L.L. Performed machine learning, model development, survival and other statistical analysis: O.I. Figures and tables: O.I. and B.G. Analyzed genotyping data: O.I., L.L. Contributed Drug Bank and genetic variant analysis tools: B.G., S.D.L., and R.C. Contributed statistical insights: H-C. L. Contributed to literature review: O.I. and N.M. Contributed to clinical interpretation: L.L., O.I. and N.M. Wrote and edited the paper: O.I., B.G., N.M., L.L., S.D.L., and J.T.D.

## Supporting information captions

**Supplementary Methods.**

**Supplementary Results.**

**Supplementary Discussion.**

**Fig. S1. Types of selected features for each disease comorbidities.** Types of selected features for each disease comorbidities. Medication, disease, lab values/vital signs, demographics were colored by blue, green, red, and purple, respectively.

**Fig. S2. Exclusion of the first 30days after T2D diagnosis.** (A) Diagnosis distribution. (B) Prescription distribution. The order of the dates of T2D and its comorbidities was important for this study, but it is not biologically meaningful to assign an exact date of diagnosis for either T2D or its comorbidities. This is especially T2D true because, from a biological point of view, T2D is subject to considerable uncertainty in regard to the date of onset. As a result, a very sharp distinction of T2D comorbidities diagnoses immediately after diagnosis of T2D is not likely to be meaningful.

**Table S1. Prevalence and increased risk after T2D.**

**Table S2. All features selected by adaptive LASSO and odds ratio estimated by logistic regression (nominal p < 0.05).**

**Table S3. All associations between T2D comorbidity and T2D /T2D-related medications (adjusted p < 0.05, using the Benjamini-Hochberg method).** References to the previous studies are also included. ^a^ Multiple published studies showed inconsistent results.

**Table S4. All protective medications identified by the survival analysis (adjusted p < 0.05, using the Benjamini-Hochberg method).**

**Table S5. All risk medications identified by the survival analysis (adjusted p < 0.05, using the Benjamini-Hochberg method)**.

**Table S6. Balance of covariates before and after propensity score matching in the survival analysis.**

**Table S7. Candidate medications for adjunctive/combination therapy with T2D medications.** (A) Significant additive effects in each subgroup of T2D patients taking a T2D medication in Table 1B. All medications in Table 1A were tested for each comorbidity, except for ‘Nutritional deficiencies’ (see Supplementary Methods). (B) Significant additive effects in each subgroup of T2D patients taking a T2D medication in Table 1A. All medications in Table 1A were tested across each comorbidity in Table 1. (C) High-resolution association between nutritional deficiency and high-risk T2D medications. Significant associations between diseases coded by ICD-9, and T2D medications (p < 0.05) clarified the type of nutritional deficiency. ^a^ Ingredient.

**Table S8. Balance of covariates before and after propensity score matching in the additive effects analysis.**

**Table S9. All association between T2D comorbidities and SNPs related to predicted protective medications (p < 0.05).**

