## Supplemental material for "Identification and genetic validation of drug-comorbidity interactions in type 2 diabetes using data-driven disease trajectory analysis"

### Supplementary Methods

#### *Patient cohort*

We utilized data from the Electronic Medical Record (EMR) system of Mount Sinai Hospital (MSH), the largest comprehensive EMR system in New York City. The system contains records for more than 7.5 million unique patients dating back to 2000. Disease diagnoses are encoded as International Classification of Diseases, 9th Revision (ICD-9) billing codes, which are often used in EMR-related analyses [1]. In this study, we retrieved records from all patients diagnosed with T2D disease (ICD-9 codes 250.x0 and 250.x2; *n*=119,847), and then performed the following pre-processing and filtering steps.

1. In compliance with Protected Health Information (PHI) and Health Insurance Portability and Accountability Act (HIPAA), we censored the ages of individuals younger than 18 or older than 90 years old to those limits.

2. We included individuals that had a healthcare visit since 2003, when the EMR was implemented in the MSH system (*n*=113,003).

3. We included individuals who did not have a history of type 1 diabetes mellitus (ICD-9 codes 250.x1 and 250.x3; *n*=103,713).

4. We included individuals with reported sex and age, and only included individuals with self-identified race/ethnicity (referred to as “race” in this manuscript) of Caucasian, African American, or Hispanic/Latino (*n*=61,275).

5.We selected patients who had at least one prescription record (n=52,183) and were followed up after more than 30 days in the EMR system (*n*=40,144).

After these filtering steps, a total of 40,144 T2D patients remained for analysis. The mean age within the population was 62.0 ± 13.1 years. The population contained 20,745 (51.7%) Males and 19,399 (48.3%) Females. The race breakdown of the population was as follows: 21,598 (53.8%) Caucasian, 13,379 (33.3%) African American, and 5,167 (12.9%) Hispanic/Latino.

Addition to patient characteristics, we retrieved all available clinical variables from EMR, including medication prescriptions, other disease diagnoses, lab values, and vital signs. In total, we compiled 5,254,956 diseases diagnosis, 7,626,493 medication prescriptions, 27,506,593 lab values, and 16,358,172 vital signs across all patients. We provide a schematic work flow of our study design and methods in Fig. 1.

##### *Clinical sources and term standardization*

We categorized diseases using the Clinical Classifications Software (CCS) for ICD-9 diagnosis codes, developed by AHRQ [2], which aggregates and characterizes more than 14,000 ICD-9 codes into broader coherent disease categories. This strategy helps to avoid sample size limitations resulting from the use of ICD-9 codes. For categorization of diseases other than neoplasms, we used the 'Single-Level CCS Diagnoses', a total of 283 different categories that classify all diagnoses into unique groups based on the ICD-9 codes. For neoplasms, we used 'Multi-Level CCS Diagnoses' to group Single-Level CCS Diagnoses into broader body systems or condition; these broader categories could enhance signals that might otherwise be lost due to the small sample size of each neoplasm subtype at the Single level. We used the root category of neoplasms, 'Neoplasms (Multi-Level CCS: 2)', and its 16 child categories

(Multi-Level CCS: 2.1-2.16), as well as a composite category we defined, 'Neoplasms excluding benign (any of Multi-Level CCS: 2.1-2.15)', i.e. any kind of neoplasm except for 'Benign Neoplasms (Multi-Level CCS: 2.16)'. We standardized medication data by mapping onto the RxNorm ontology [3]. Specifically, we mapped these terms to ingredient-level codes, and unified all types of insulins into a single code, yielding 2,928 normalized medications. We standardized lab value and vital sign data by mapping onto LOINC ontology codes [4]. For each lab value and vital sign, the value was summarized as normal (i.e., within the normal range), higher than the normal range, or lower than the normal range.

##### *Definition of outcomes and observation period of covariates in the machine-learning model*

We defined "T2D medications" based on eMERGE definition [5] and added newly FDA approved medications, including DPP-4 inhibitors (vildagliptin, saxagliptin, alogliptin, and linagliptin), GLP-1 agonists (liraglutide, lixisenatide, albiglutide, and dulaglutide) and SGLT-2 inhibitors (dapagliflozin, canagliflozin, and empagliflozin).

We defined "T2D-related medications" as medications known to be useful for treating or managing T2D, excluding T2D medications, according to knowledgebase MEDI high-precision subset (MEDI-HPS) with 'possible label use' marker [6]. T2D-related medications include statins to treat lipid disorders and antihypertensive medications, including angiotensin II receptor (*AT2R*) blockers and angiotensin-converting-enzyme (*ACE*) inhibitors.

We defined the initial T2D diagnosis date as the day of first T2D diagnosis or the first prescription day of T2D medications, whichever occurred earlier. In our EMR system, there are 13% T2D patients (n=6,950/52,183) who have a record of prescription of T2D medication before the ICD-9 code–based diagnosis of T2D (median=8 days, inter quartile range (IQR) = 2–88 days), which is a relatively short delay in comparison with comorbidity onset.

We used Single-Level CCS Diagnoses terms to define outcome comorbidities except for neoplasms and Multi-level CCS Diagnoses for neoplasms as described above. We selected the T2D comorbidities based on the disease prevalence analysis detailed below. To discover risk factors or new therapeutic options for T2D sequelae, we focused on new onset of comorbidities appearing more than 30 days after the initial T2D diagnosis date in order to avoid uncertainty in the sequence of T2D and the comorbidity [7] (Supplementary Fig. S2). We collected a full history of diagnosed diseases, lab tests, and vital signs before the first onset of comorbidities. We also retrieved prescribed medications from one year prior to the initial outcome diagnosis date to the first onset of T2D comorbidities.

##### *Disease prevalence*

We calculated the disease prevalence based on the Kaplan–Meier product–limit estimator, with units of 1 year.

The Kaplan–Meier product–limit estimator is defined as

$$S(t_i) = \prod_{t_i \leq t} \frac{N_{year}^{i} - D_{year}^{i}}{N_{year}^{i}} \qquad \text{(S1)}$$

where $S(t_i)$ is the estimated survival probability for any particular one of *t* time periods, $N_{year}^{i}$ is the number of individuals at risk at the beginning of year $i$, and $D_{year}^{i}$ is the number of events during time period $t_i$. Disease prevalence is $1 - S(t_i)$.

We selected T2D comorbidities for which prevalence within 10 years after the initial T2D diagnosis was greater than 10%, or which belonged to any of the neoplasm categories. In total, 109 comorbid diseases were used as T2D comorbidities in the following analyses.

##### *Disease pair temporal directionality*

For all patients with T2D, we assessed disease-pair connectivity patterns for T2D comorbidities. Specifically, we determined whether the members of each pair exhibited a significant pattern in their temporal ordering, e.g., whether one preceded the other more often than expected by chance. We performed a cumulative binomial probability test to assess the temporal ordering of the associations between T2D and each of the T2D comorbidities, giving a 50% probability of T2D to occur before the comorbid disease.

##### *Logistic regression model*

We used the odds ratio (OR) from logistic regression (Eq. S2) to quantify the magnitude of the risk of comorbidity associated with prescription of each medication (i.e., increased risk or protective effect) after adjusting for clinical confounders selected by adaptive LASSO, patient demographics, and follow-up time frame. In addition, we always adjusted for insulin prescription, because insulin intervention affects the condition of T2D patients and is likely to reflect an advanced disease state [8].

$$log\left(\frac{P}{1-P}\right) = \beta_0 + \sum_x \beta_x LASSO\_selected\_covariate_x + \beta_i insulin + \beta_a age + \beta_g gender + \beta_r race + \beta_p observed\ period \qquad \text{(S2)}$$

where $P$ is the probability of a disease, $LASSO\_selected\_covariate_x$ are the adaptive-LASSO-selected covariates, including medications, disease history, and lab/vital values, is a binary variables, *insulin* is a binary variable, *age* is a continuous parameter, *gender* is a binary variable (Female/Male), *race* is a categorical variable (Caucasian, African American, or Hispanic/Latino), and observed period is a continuous parameter. $\beta$ coefficients for each covariate represent the effect size when controlling for all others.

Features that were significantly associated with each of the outcome comorbid diseases with nominal p values less than 0.05 were used for the following propensity score matching and survival analysis.

##### ***Propensity score matching***

To control for potential confounding factors due to imbalances of clinical characteristics, such as demographics, we analyzed the temporal effects of other covariates, such as medications, diseases, lab values and vital signs after propensity score matching (using the R package *MatchIt*) [9] to select the matched control cohort for the targeted case cohort [10]. In this manner, we created comparable cohorts, consisting of groups treated or not treated with a targeted medication, based on a set of covariates at the baseline time point (Fig. 1B). The baseline time point was defined as the first prescription day of the targeted medication or 30 days after the initial T2D diagnosis date, whichever was later, because we observed disease comorbidities for more than 30 days after the initial T2D diagnosis date in our hospital [7].

The propensity scores of targeted prescriptions were predicted by a logistic regression model, including other significant confounders selected by adaptive LASSO with patient demographics as covariates. Each patient prescribed a given medication was matched to a corresponding comparison patient within a 0.2 caliper of propensity score by nearest-neighbor matching. Patients in both treated and control groups outside the support of the distance measure were discarded [9].

We set the matching ratios flexibly to maximize the number of control patients to match to each treated patients [11], because the number of treated patients varied widely (maximum was 20,688 patients for acetaminophen; minimum was 22 patients for ilohexol and minocycline). The matched mean distance between treated and control groups revealed that the matched cohort was well balanced (median: 0.0043, Inter quartile range [IQR]: 0.0019-0.011, maximum: 0.064) (Supplementary Table S6).

##### *Additive effects of two medications*

We investigated whether two medications in combination could produce a sum effect based on their individual effects, i.e. an additive effect. For this purpose, we selected each subgroup of patients who were on T2D medications, and then we calculated the hazard ratios of the second medications by performing a survival analysis using propensity score matching and time-dependent covariates (Supplementary Table S8), except for the definition of the baseline time. We set the baseline time point as the first prescription day of the T2D medication or another medication, whichever occurred later. We analyzed the combinations of the T2D medications significantly associated with the risk of the comorbidities and the significantly protective medications. In addition, we tested the combinations of two significantly protective medications in the same manner. We analyzed six well-known T2D complications associated with T2D medications, including two major microvascular complications, ʻRetinal detachments; defects; vascular occlusion; and retinopathyʼ and ʻChronic kidney diseaseʼ; two major macrovascular complications ʻCoronary atherosclerosis and other heart diseaseʼ and ʻAcute cerebrovascular diseaseʼ; and two comorbidities that were enriched in particular T2D subtypes in our previously published study [12], ʻNeoplasm excl. benignʼ and ʻDelirium, dementia, and amnestic and other cognitive disordersʼ. In addition to these six comorbidities, ʻNutritional deficiencyʼ is also well-known T2D complication and was enriched in a T2D subtype [12]. We clarified the type of the nutrient in the subsequent high-resolution association study, because the direct and effective intervention would be adequate intake of the appropriate nutrient.

##### *High-resolution association study between nutritional deficiency and medications*

To nail down the specific type of nutritional deficiency, we studied the association between a medication and each disease coded by ICD-9 (260.x-269.x and 799.4), which comprise the 'Nutritional deficiency' Single-Level CCS diagnoses. In particular, we analyzed four T2D medications related to the risk of 'Nutritional deficiency', namely metformin, glyburide, repaglinide, and nateglinide. Odds ratios were calculated by Fischer's exact test by counting the patients who were or were not on the medication, and who were diagnosed with one of the ICD-9 codes listed above.

##### *Biobank cohort*

As of 2016, the Charles Bronfman Institute of Personalized Medicine Bio*Me* biobank (http://icahn.mssm.edu/research/ipm) within the Icahn School of Medicine at Mount Sinai has collected genetic data for over 30,000 patients with linked EMR [13]. A subset of Bio*Me*, consisting of over 11,000 individuals, were genotyped using the Illumina Human Omni Express Exome Bead-8 BeadChip v1.1 array. We selected T2D patients using the same criteria as described above for the EMR (i.e., filtering steps 1–3), yielding 2,808 patients. The mean age was 61.3 ± 12.7 years. This cohort consisted of 1,110 (39.5%) Males and 1,698 (60.5%) Females. The race breakdown was as follows: 341 (12.1%) Caucasian, 982 (35.0%) African American, 1,434 (51.1%) Hispanic/Latino, and 51 (1.8%) Other.

##### *SNP/gene and disease association analyses*

We assessed our predicted associations between medications and above seven well-known T2D comorbidities [12] by performing a genetic association analysis. Specifically, we retrieved drug–gene relationship information from DrugBank version 5.0.11 [14], and then obtained genotyping data for all SNPs mapped to these genes in our cohort. We analyzed association aSNP–comorbidity basis as well as a gene–comorbidity basis.

We performed logistic regression for the SNP–comorbidity combinations. We also controlled for patient demographics, including age, self-reported sex, and genetic ancestry using Principal Component Analysis (PCA) with the first five principal components, which explained the majority of variance (Eq. S3). The use of PCA on genetic data for determining and controlling for genetic ancestry in association studies is well established [15]. We focused on the significance of the SNP and the magnitude and direction of the associated β value, which represents the effect size after controlling for other covariates (positive values indicate increased risk).

$$log\left(\frac{P}{1-P}\right) = \beta_0 + \beta_s SNP + \beta_a age + \beta_g gender + \sum_{i=1}^{5} \beta_i \cdot PC_i \quad \text{(S3)}$$

where *P* is the probability of a disease, *SNP* is a binary variable, age is a continuous parameter, *gender* is a binary variable (Female/Male), and $PC_1$–$PC_5$ are continuous parameters. $\beta$ coefficients for each covariate represent effect size when controlling for all others.

Additionally, we analyzed genetic associations at the gene level by using the SNP-set Kernel Association Test (SKAT) method [16], which uses a kernel machine to tests for association between an outcome and both rare and common variants. SKAT aggregates data for all measured variants within a region (in this case, a gene) and assesses the association between the outcome and this variant burden. We controlled for all covariates used in the SNP-level analysis. We only evaluated the 149 genes that were drug targets for any of 40 medications ($n$=4,912 for the SNP-level analysis and $n$=172 for the gene-level analysis across seven comorbidities). As the purpose of this analysis was for validation, and not for discovery study, we used a nominal P-value threshold for significance. We implemented SKAT using the R package *SKAT*.

### Supplementary Results

##### *The increase in risk of each comorbidity after T2D diagnosis*

The increase in risk of each comorbidity after T2D diagnosis was quantified based on the timing of their occurrence in individual patients. Specifically, for each comorbid disease pair (i.e. T2D and another disease category), we tabulated the number of patients with both diseases and assessed which of the two diseases occurred first. Overall, out of the 109 T2D comorbidities analyzed, 93 diseases were significantly more likely to occur after T2D diagnosis (adjusted $p < 0.05$; Fig. 2B and Supplementary Table S1). We identified some disease associations previously reported to be specifically enriched in T2D subtypes [17], including microvascular complications such as 'Retinal detachments; defects; vascular occlusion; and retinopathy' [Relative risk (RR): 1.76, adjusted p <0.0001], 'Acute and unspecified renal failure' (RR: 1.69, adjusted $p < 0.0001$), and 'Delirium, dementia, and amnestic and other cognitive disorders' (RR: 1.64, adjusted $p < 0.0001$), as well as 'Nutritional deficiencies' (RR: 1.70, adjusted $p < 0.0001$).

##### *Selection of features for each T2D comorbidity by cross-sectional analysis*

Our first task for medication-comorbidity association was to identify the important features for each T2D comorbidity, specifically medications (N=899), lab/vital signs (N=107), disease diagnoses (N=246), and patient demographics (N=4) prior to the onset of each comorbidity. A total of 551 unique medications were selected as significant features associated with at least one T2D comorbidity (Supplementary Fig. S1 and Supplementary Table S2). On average, 83.7 ± 28.5 (mean ± SD) features were selected for each comorbidity: 37.0 ± 15.2 medications, 28.9 ± 10.2 disease diagnosis, 15.3 ± 5.5 lab/vital signs, and 2.6 ± 1.1 demographic characteristics.

Among the 551 unique medications selected as significant features, 331 had a total of 1,726 protective associations, and 452 had a total of 2,228 risk associations (nominal $p < 0.05$; Supplementary Table S2) with T2D comorbidities. Two hundred and thirty-two medications had both protective and risk associations with T2D comorbidities, suggesting that they decrease the risk of some diseases but increase the risk of others.

##### *Candidate medications for adjunctive/combination therapy with T2D medications*

We explored candidate medications for adjunctive/combination therapy in order to devise safe and effective therapies capable of reducing complication probabilities. For this purpose, we selected each subgroup of T2D patients taking one of medications in Tables 1A or 1B, and we analyzed the association between other medications in Table 1A and comorbidities. The results of these subgroups' analysis are shown in Supplementary Tables S7A and S7B.

Seven medications were associated with decreased risk for 'Retinal detachments; defects; vascular occlusion; and retinopathy', including clopidogrel (HR=0.77, p=0.0094), ibuprofen (HR=0.69, p=0.0021), calcium (HR=0.61, p=0.0019), and psyllium (HR=0.26, p=0.0034) not only in entire T2D cohort but also in the subgroup of insulin users (Supplementary Table S7A). Insulin users taking metformin were likely to decrease the risks of 'Chronic kidney disease' and 'Coronary atherosclerosis and other heart disease' (HR=0.64, p<0.0001; HR=0.68, p<0.0001, respectively). Furthermore, we found six non-T2D-related medications, including acetaminophen (HR=0.63, <0.0001) and ondansetron (HR=0.70, p<0.0001), that may protect against for 'Chronic kidney disease' in patients using insulin (Table 2A). Six other non-T2D-related medications, including cholecalciferol (HR=0.55, p=0.0050) and iron (HR=0.69, p=0.0083) could prevent 'Coronary atherosclerosis and other heart disease' in patients using insulin (Supplementary Table S7A).

Metformin, consistently with previous studies, exhibited protective effects against 'Coronary atherosclerosis and other heart disease,' 'Delirium, dementia, and amnestic and other cognitive disorders,' and 'Chronic kidney disease' (Table 1A). Our analysis revealed nine medications, which both reduced these risks individually, and also had an additive protective effect on metformin users (Supplementary Table S7B). These medications include ondansetron and naproxen for 'Chronic kidney disease' (HR=0.69, p=0.0020 and HR=0.65, p=0.014, respectively), pseudoephedrine and iron for 'Coronary atherosclerosis and other heart disease' (HR=0.46, p=0.0080 and HR=0.57, p=0.0026, respectively), and five medications (atorvastatin, gabapentin, ondansetron, hydromorphone, and phenylephrine) for 'Delirium, dementia, and amnestic and other cognitive disorders'.

We narrowed down the association between four T2D medications and risk of 'Nutritional deficiencies,' as coded by ICD-9 (260.x-269.x and 799.4), which supports monitoring and use of adjunctive supplements where indicated (Supplementary Table S7C). Metformin was significantly associated with 'other B-complex deficiencies (266.2)' (OR=1.36, p=0.00078), which is consistent with published studies [18], and also with 'unspecified vitamin D deficiency' (ICD-9 code: 268.9) (OR=1.57, p<0.0001), which has not previously been described. Other novel associations included a sulfonylurea, glyburide related to 'other B-complex deficiencies' (OR=1.57, p=0.035). Repaglinide, a meglitinide, was associated with 'unspecified protein-calorie malnutrition (263.9)' (OR=1.75, p=0.0082), and another meglitinide, nateglinide, was linked to 'unspecified vitamin D deficiency' (OR=2.21, p=0.018).

### Supplementary Discussion

##### *Antibiotics and neoplasms associations supported by the previous laboratory and animal studies*

The antibiotic bacitracin exhibited a protective effect against ʻNeoplasm excl. benignʼ, especially ʻSecondary malignanciesʼ (Supplementary Table S4). A recent experiment based on a cell-based assay showed that bacitracin inhibited the migration of glioma cells by interfering with the integrin pathway, which plays a major role in cell migration and invasion [19]. Because secondary malignancies occur due to migration of the primary cancer, our clinical findings are supported by this cell-based study. Another antibiotic, azithromycin, also showed protective effect against ʻNeoplasm excl. benignʼ. This association is consistent with a published study showing that the inhibitory effects of azithromycin on angiogenesis and lung tumor growth in mouse models are mediated by suppression of *VEGF* receptor 2 [20]. Moreover, we discovered a novel protective association between vancomycin and ʻNeoplasms excl. Benignʼ and ʻOther gastrointestinal cancerʼ. This finding is interesting because T2D is associated with an increased overall cancer risk and an increased risk of gastrointestinal cancers in particular [21,22].

##### *Additional potentially novel association supported by genetic analysis*

The association between ‘Delirium, dementia, and amnestic and other cognitive disorders’ and the antibiotic cefazolin was linked to an unintended target of cefazolin, *PON1*. PON1 binds to high-density lipoprotein (HDL) particles, and its expression is differentially regulated in *APOE3* and *APOE4* transgenic mice; *APOE* genotype is the primary genetic risk factor for Alzheimer’s disease [23]. Although cefazolin is not likely to penetrate the blood–brain barrier [24], we hypothesize that it might affect cognitive disorders by perturbing lipid turnover or tissue distribution.

Another potentially novel protective association between 'Acute cerebrovascular disease' and repaglinide, belonging to the meglitinide class of T2D medications, was validated by the relationship between this disease and main therapeutic targets of repaglinide: *ABCC8*, which encodes sulfonylurea receptor 1 (SUR1), and *PPARG*, which encodes peroxisome proliferator-activated receptor gamma (*PPARG*). In addition to the indirect benefits of lowering glucose levels in the blood via its action on these proteins, the protective effect of repaglinide could be explained by neurodegenerative and inflammatory process in the brain via the same proteins. In rodent models of stroke, *SUR1* is expressed in reactive microglial cells, and blockage of *SUR1* improves outcomes by preventing brain swelling and enhancing neuroprotection [25,26]. Furthermore, the activation of intracerebral *PPARG* protects against cerebral ischemia in rodents by various mechanisms, including antioxidative activity, suppression of neurodegenerative target genes, and inhibition of inflammation [27].

The relationship between 'Retinal detachments; defects; vascular occlusion; and retinopathy' and the NSAID ibuprofen was supported by 27 SNPs in 7 genes, including *PTGS2* (encoding cyclooxygenase 2) and *PPARA* and *PPARG* (encoding peroxisome proliferator–activated receptor). In a diabetic rat model, expression of *PPARA,* but not *PPARG,* is significantly down-regulated in the retina, and diabetic *PPARA* KO mice develop more severe diabetic retinopathy [28]. Because ibuprofen activates both *PPARA* and *PPARG* [29], its protective effects against retinopathy might be mediated via its effect on *PPARA*, in addition to the anti-inflammatory effect mediated by cyclooxygenase 2.

##### *Specific treatment strategies according to individual T2D disease*

In our current study, we build upon our previous work to identify specific treatment strategies according to individual T2D disease profiles and drug-disease associations. In our preceding publication, we characterized the heterogeneous patient landscape of T2D into three distinct subtypes using clinical data from an EMR and verified the disease association in a genetics biobank [12]. That work had suggested that there might be patterns of T2D with different genetic associated phenotypic characteristics, as well as risk factors for developing certain subsequent conditions. Combining this knowledge with our current observations regarding medication associations with comorbidities, a more personalized and precise approach toward medication regimens can be devised. For example, adjunctive treatment with clopidogrel and/or NSAIDs, such as ibuprofen, could be useful for preventing retinopathy in T2D subtype 1, which is characterized by diabetic retinopathy. For T2D subtype 2, enriched for cancer malignancy and cardiovascular disease, universal prophylactic administration of low-dose aspirin could be considered, given that it was demonstrated to have a protective association with lung cancer (Supplementary Table S4) and that it reduces the incidence of adverse cardiovascular events and all-cause mortality, in patients with established cardiovascular disease [30]. For subtype 3, associated with cardiovascular and neurological diseases, a greater emphasis can be placed on metformin and statin use, because these are associated with protective effects on coronary atherosclerosis and other heart disease, as well as cognitive disorders. Furthermore, the DPP-4 inhibitor sitagliptin could decrease the future risk of acute cerebrovascular disease. In the examples delineated above, one can perceive how results from our combined studies can pave the way for tailoring treatment regimens based on personal risk profiles.

### Supplementary Figures

Supplementary Fig. S1

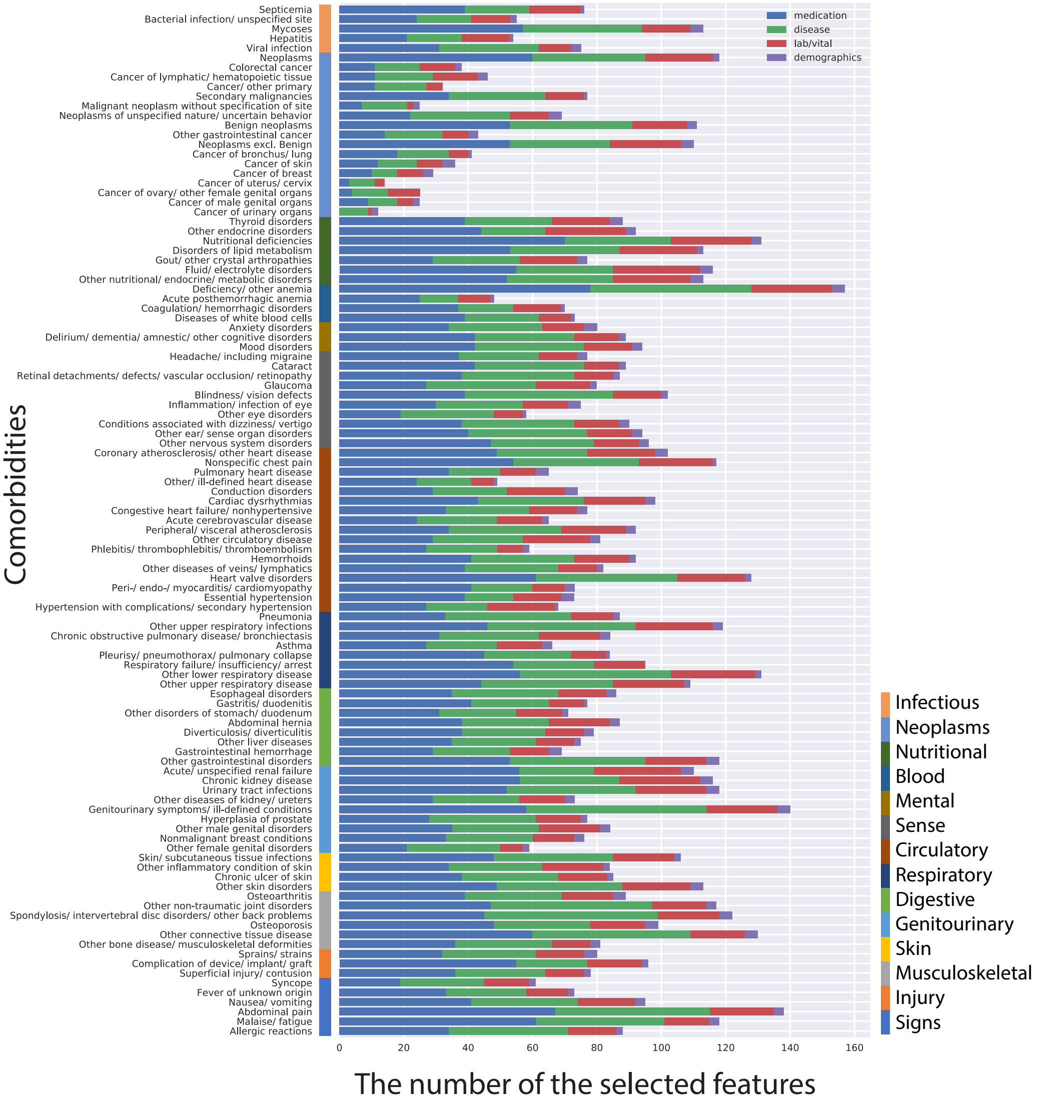


**Fig. S1. Types of selected features for each disease comorbidities.** Types of selected features for each disease comorbidities. Medication, disease, lab values/vital signs, demographics were colored by blue, green, red, and purple, respectively.

Supplementary Fig. S2

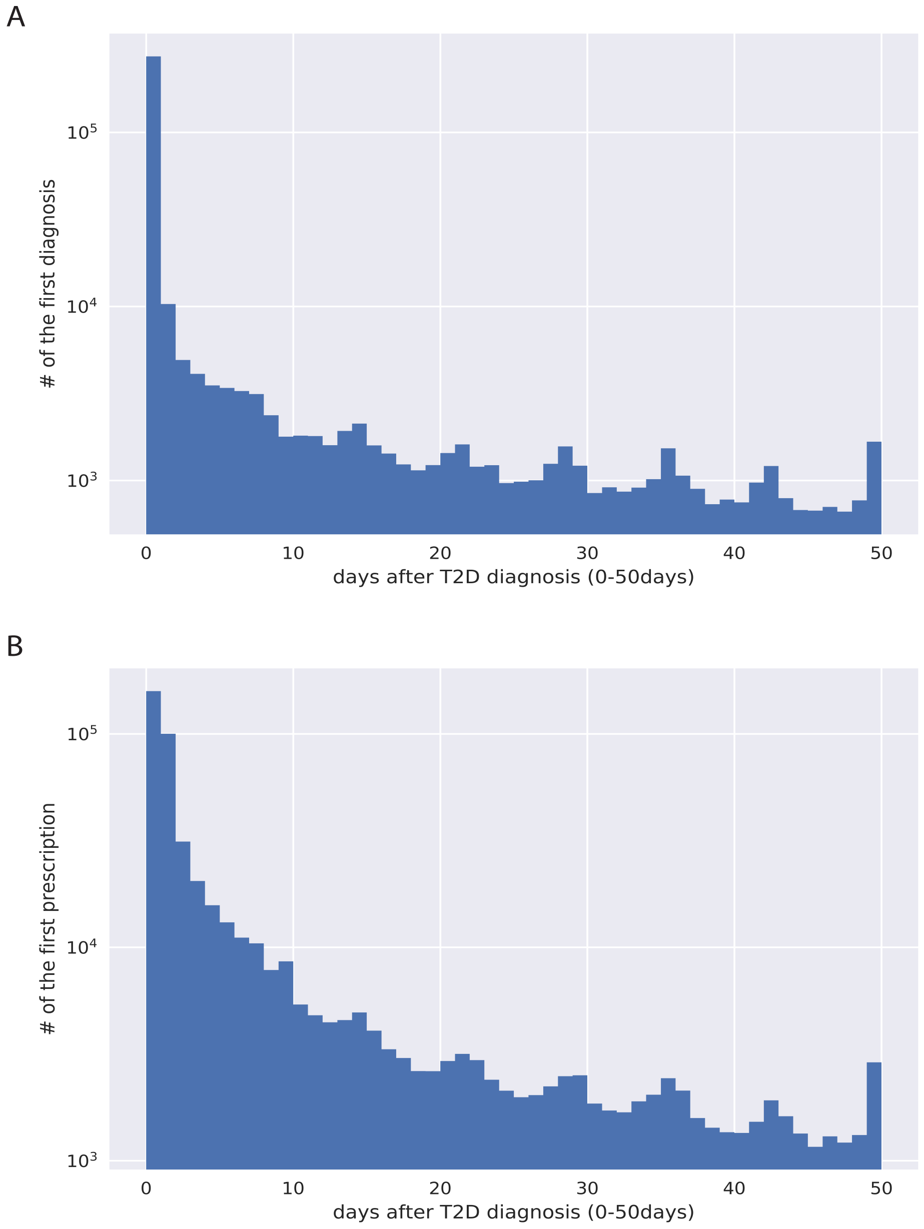


**Fig. S2. Exclusion of the first 30days after T2D diagnosis.** (A) Diagnosis distribution. (B) Prescription distribution. The order of the dates of T2D and its comorbidities was important for this study, but it is not biologically meaningful to assign an exact date of diagnosis for either T2D or its comorbidities. This is especially T2D true because, from a biological point of view, T2D is subject to considerable uncertainty in regard to the date of onset. As a result, a very sharp distinction of T2D comorbidities diagnoses immediately after diagnosis of T2D is not likely to be meaningful.

### Supplementary tables (Excel files)

**Table S1. Prevalence and increased risk after T2D.**

**Table S2. All features selected by adaptive LASSO and odds ratio estimated by logistic regression (nominal p < 0.05).**

**Table S3. All associations between T2D comorbidity and T2D /T2D-related medications (adjusted p < 0.05, using the Benjamini-Hochberg method).** References to the previous studies are also included. [a] Multiple published studies showed inconsistent results.

**Table S4. All protective medications identified by the survival analysis (adjusted p < 0.05, using the Benjamini-Hochberg method).**

**Table S5. All risk medications identified by the survival analysis (adjusted p < 0.05, using the Benjamini-Hochberg method).**

**Table S6. Balance of covariates before and after propensity score matching in the survival analysis.**

**Table S7. Candidate medications for adjunctive/combination therapy with T2D medications.** (A) Significant additive effects in each subgroup of T2D patients taking a T2D medication in Table 1B. All medications in Table 1A were tested for each comorbidity, except for 'Nutritional deficiencies' (see Methods). (B) Significant additive effects in each subgroup of T2D patients taking a T2D medication in Table 1A. All medications in Table 1A were tested across each comorbidity in Table 1. (C) High-resolution association between nutritional deficiency and high-risk T2D medications. Significant associations between diseases coded by ICD-9, and T2D medications (p < 0.05) clarified the type of nutritional deficiency. [a] Ingredient.

**Table S8. Balance of covariates before and after propensity score matching in the additive effects analysis.**

**Table S9. All association between T2D comorbidities and SNPs related to predicted protective medications ($p < 0.05$).**
